# Predicting RNA 3D structure and conformers using a pre-trained secondary structure model and structure-aware attention

**DOI:** 10.1101/2025.04.09.647915

**Authors:** Wenkai Wang, Zhenling Peng, Jianyi Yang

## Abstract

Determining RNA 3D structure and conformers remains a grand challenge in structural biology. In this work, we propose trRosettaRNA2, a deep learning-based end-to-end approach to this problem. Considering the scarcity of RNA 3D structure data, trRosettaRNA2 integrates an auxiliary secondary structure (SS) prior module, pre-trained on extensive SS data, to generate informative base-pairing priors. This module also serves as an independent RNA secondary structure prediction method, trRNA2-SS, and achieves state-of-the-art performance. To enable end-to-end prediction, trRosettaRNA2 employs SS-aware attention to generate RNA 3D structure and conformers. Rigorous benchmarks demonstrate that trRosettaRNA2 outperforms other RNA 3D structure prediction methods, despite using significantly fewer parameters and computational resources. Notably, its flexibility in leveraging diverse secondary structure inputs provides a pathway to generate accurate 3D structure and explore the RNA conformers. Based on trRosettaRNA2, our group, Yang-Server, ranks as the top automated server for RNA structure prediction in the CASP16 blind test, surpassing AlphaFold3. This performance highlights that trRosettaRNA2 represents a solid step towards the “AlphaFold moment” for RNA structure prediction. Application to the RNase P RNA demonstrates that trRosettaRNA2 successfully captures its structural heterogeneity even without requiring experimental data, showing its potential to predict RNA conformational ensembles.

## INTRODUCTION

Determining the structures and conformations of the Ribonucleic acids (RNAs) is crucial for elucidating their biological functions, such as protein synthesis, gene regulation, and ribozyme catalysis, which are fundamental to life. However, due to their inherent flexibility and structural instability, solving RNA 3D structures and conformations with experimental techniques like X-ray crystallography, Nuclear Magnetic Resonance (NMR) spectroscopy, and cryogenic electron microscopy (cryo-EM) is significantly more challenging than for proteins ^1^. To address this challenge, considerable effort has been dedicated to developing computational methods for RNA structure prediction over the last decades. These methods are primarily based on fragment assembly, template-based modeling, and/or physical-based modeling ^2–8^. While these methods have shown promise in modeling certain small RNAs, their accuracy and computational cost remain major obstacles when dealing with the broader spectrum of RNA challenges.

Inspired by the breakthrough of deep learning in protein structure prediction ^9–12^, several deep learning-based methods have been proposed to tackle the RNA structure prediction challenges in recent years ^13–20^. For example, DeepFoldRNA ^14^ uses a transformer neural network to predict the inter-residue geometry from multiple sequence alignment (MSA) and secondary structure (SS), subsequently folding the 3D structure through energy minimization. RhoFold+ ^16^ introduces an RNA language model to extract additional co-evolutionary signals that complement MSA information. RoseTTAFoldNA ^18^ employs an end-to-end neural network to predict the 3D structures of RNA, DNA, and nucleic acid-protein complexes. RoseTTAFold All-Atom ^19^ and AlphaFold 3 ^20^ were proposed to predict structures for all biomolecules, including RNA. trRosettaRNA ^13^, our previous method, adopts a Res2Net ^21^-enhanced transformer architecture (RNAformer) to improve 2D geometry prediction. According to the benchmark tests and blind tests in CASP15 and RNA-Puzzles, trRosettaRNA significantly outperforms traditional automated approaches and achieves competitive performance with human experts for structure prediction of natural RNAs.

Nevertheless, the CASP15 competition also highlighted that deep learning has not yet achieved the “AlphaFold moment” for RNA ^22, 23^. The most evident examples are the four synthetic RNAs from CASP15, for which all automated methods failed to reach performance comparable to human experts. The challenge stems from two primary aspects. First, the limited amount of experimentally determined RNA 3D structure data available in the Protein Data Bank (PDB) ^24^ (only ∼4% entries containing RNAs) hinders the performance of data-driven methods. Second, the relatively simple chemical alphabet of RNA (four bases) compared to protein (20 amino acids), and the limitation of current alignment algorithms, lower the quality of RNA MSAs ^23^, which is essential in the paradigm inherited from protein structure prediction.

Recently, a few studies have begun to predict RNA conformations by integrating computational methods with experimental data. For example, HORNET, a deep neural network, was proposed to determine RNA 3D conformations from atomic force microscopy (AFM) images ^25^. This approach successfully determined the heterogeneous conformations of ribonuclease P (RNase P) RNAs ^26^. However, modeling the inherent heterogeneity of RNA structures without relying on experimental data remains a major challenge.

The limited experimental 3D structure data and the high heterogeneity of RNA molecules pose significant challenges to RNA biology. It is widely believed that RNA 3D structure folds after forming the base-paring interactions (i.e., 2D secondary structure) ^27–29^. The conformational heterogeneity may also be inferred from its secondary structures. Unlike RNA 3D structures, RNA secondary structures are easier to experimentally determine, especially using high-throughput chemical probing techniques such as SHAPE ^30^ and DMS ^31^. For instance, 102,318 secondary structures are available in the bpRNA database ^32^, ∼12 times the number of the RNA 3D structures in PDB. Therefore, we argue that an effective solution to the prediction of RNA 3D structures and conformers can come from a close connection between RNA 2D and 3D structures. Specifically, we sought to develop a novel secondary structure prediction module, which can be used to guide the subsequent generation of 3D structure and conformers.

Following this idea, we propose trRosettaRNA2, a new end-to-end method to predict RNA 3D structure and conformers. To address the data scarcity problem for RNA 3D structures, trRosettaRNA2 incorporates a secondary structure (SS) module, which was pre-trained on a large SS dataset and can provide prior information about base-pairing interactions. Evaluated as an independent secondary structure predictor, it achieves state-of-the-art performance. To facilitate end-to-end prediction, we introduce an SS-aware structure module to directly predict the 3D structure. Benchmark tests suggest that trRosettaRNA2 outperforms other methods. Leveraging trRosettaRNA2, our group, Yang-Server, ranked as the top automated server in CASP16. Finally, trRosettaRNA2 successfully generates conformations matching well with the experimental data for RNase P RNA solved by AFM.

## RESULTS

### Overview of trRosettaRNA2

As depicted in Fig. 1, trRosettaRNA2 employs an end-to-end pipeline for RNA 3D structure prediction. Beginning with the nucleotide sequence of the target RNA, a multiple sequence alignment (MSA) is first generated by searching the RNAcentral database ^33^ using Infernal ^34^, which is then embedded into a numerical representation. A secondary structure (SS) module is designed to enhance model performance by providing prior knowledge of base-pairing. This module was pre-trained on the bpRNA secondary structure database ^32^, allowing it to effectively leverage the rich information within the extensive secondary structure data and overcome the data scarcity in RNA 3D structure prediction. Notably, this SS prior module can also function as an independent RNA secondary structure predictor, named trRNA2-SS.

**Fig. 1.**
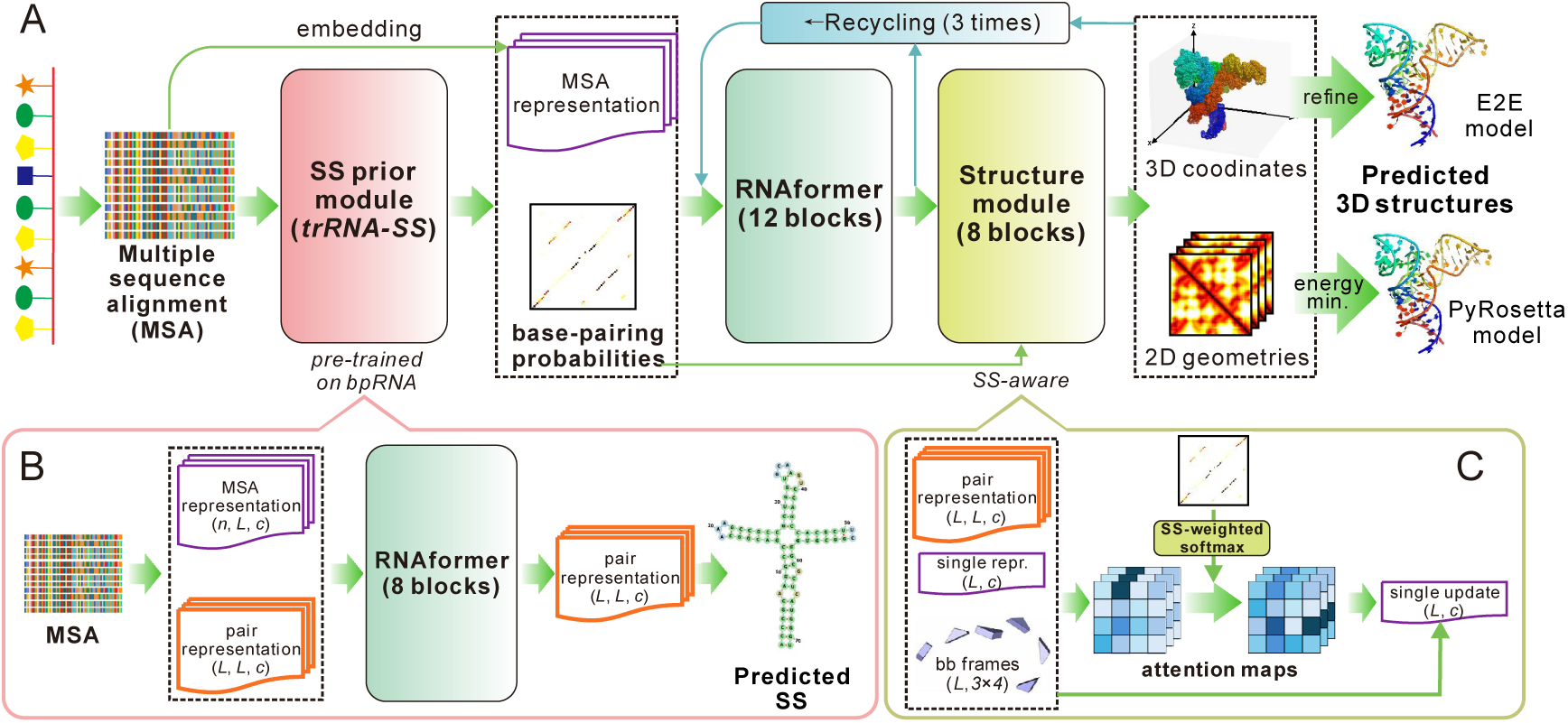
Overview of trRosettaRNA2 pipeline. (A) Pipeline for RNA 3D structure prediction using trRosettaRNA2. (B) Architecture of the SS prior module (i.e., the pipeline for RNA secondary structure prediction with trRNA2-SS). (C) Overall architecture of an SS-aware IPA block.

The output of the SS prior module (in the form of the base-pairing probabilities), alongside the MSA embedding, is then fed into a modified RNAformer module to interactively update the 1D and 2D representations of the RNA. Finally, an SS-aware structure module is employed to predict the Cartesian coordinates of all heavy atoms, which are then quickly relaxed within PyRosetta ^35^ to reduce structural violations. Alternatively, if there are severe clashes in the E2E model, deep learning constraints-guided energy minimization is used to generate more physicochemically plausible structures. These two approaches are termed trRosettaRNA2 (E2E) and trRosettaRNA2 (PyRosetta), respectively. Unless otherwise specified, “trRosettaRNA2” refers to the trRosettaRNA2 (PyRosetta) method.

### Performance of trRNA2-SS for RNA secondary structure prediction

The limited data of experimentally determined 3D RNA structures is a primary bottleneck hindering the generalization of deep learning methods in RNA structure prediction. To overcome this, we introduce an SS prior module (Fig. 1B), which was pre-trained on the extensive secondary structure data from bpRNA ^32^, to infuse 2D prior knowledge into the 3D structure prediction process. Meanwhile, this module also serves as an independent RNA secondary structure predictor, named trRNA2-SS. trRNA2-SS is trained using a transfer learning approach: initial pre-training on the bpRNA dataset, followed by fine-tuning on a smaller dataset derived from PDB (see Methods).

To rigorously evaluate the performance of trRNA2-SS, we collected two independent test sets. The first test set includes 279 RNAs from ArchiveII ^36^, a widely used SS benchmark set. The second set, PDB27, includes 27 non-redundant RNA-only entries from PDB, all released after 2022-01. Both datasets were rigorously filtered to eliminate redundancy with our training data (see Methods for details).

We compare trRNA2-SS against a serial of representative methods, including traditional energy- and/or evolution-based approaches (EternaFold ^37^, RNAfold ^38^, RNAstructure ^39^, PETfold ^40^) and deep learning-based methods (RNA-FM ^41^, RNA-MSM ^42^, UFold ^43^, MXfold2 ^44^, SPOT-RNA ^45^, SPOT-RNA2 ^46^). Among them, PETfold, SPOT-RNA2, and RNA-MSM are based on MSA, whereas the other methods rely solely on the single sequence.

As shown in Fig. 2, trRNA2-SS outperforms all compared methods across both test sets, in terms of both global and per-target AUPRC. Notably, the PDB27 dataset presents a greater modeling difficulty than ArchiveII, as indicated by the lower precision-recall curve (Fig. 2B vs. Fig. 2A) and the decreased per-target AUPRC values (Fig. 2C). For example, the AUPRC values of RNA-MSM and PETfold drop significantly from >0.8 on the ArchiveII set to <0.5 on the PDB27 set. We found that ∼95% (265/279) of RNAs in the ArchiveII set belong to at least one Rfam family (based on Infernal searches against the Rfam 14.7 database ^47^). In contrast, only 22.2% (6/27) of RNAs in the PDB27 set are associated with Rfam families. This difference explains the greater difficulty of the PDB27 RNAs, especially for RNA-MSM and PETfold, which were trained on Rfam datasets filtered to exclude families with experimentally determined structures.

**Fig. 2.**
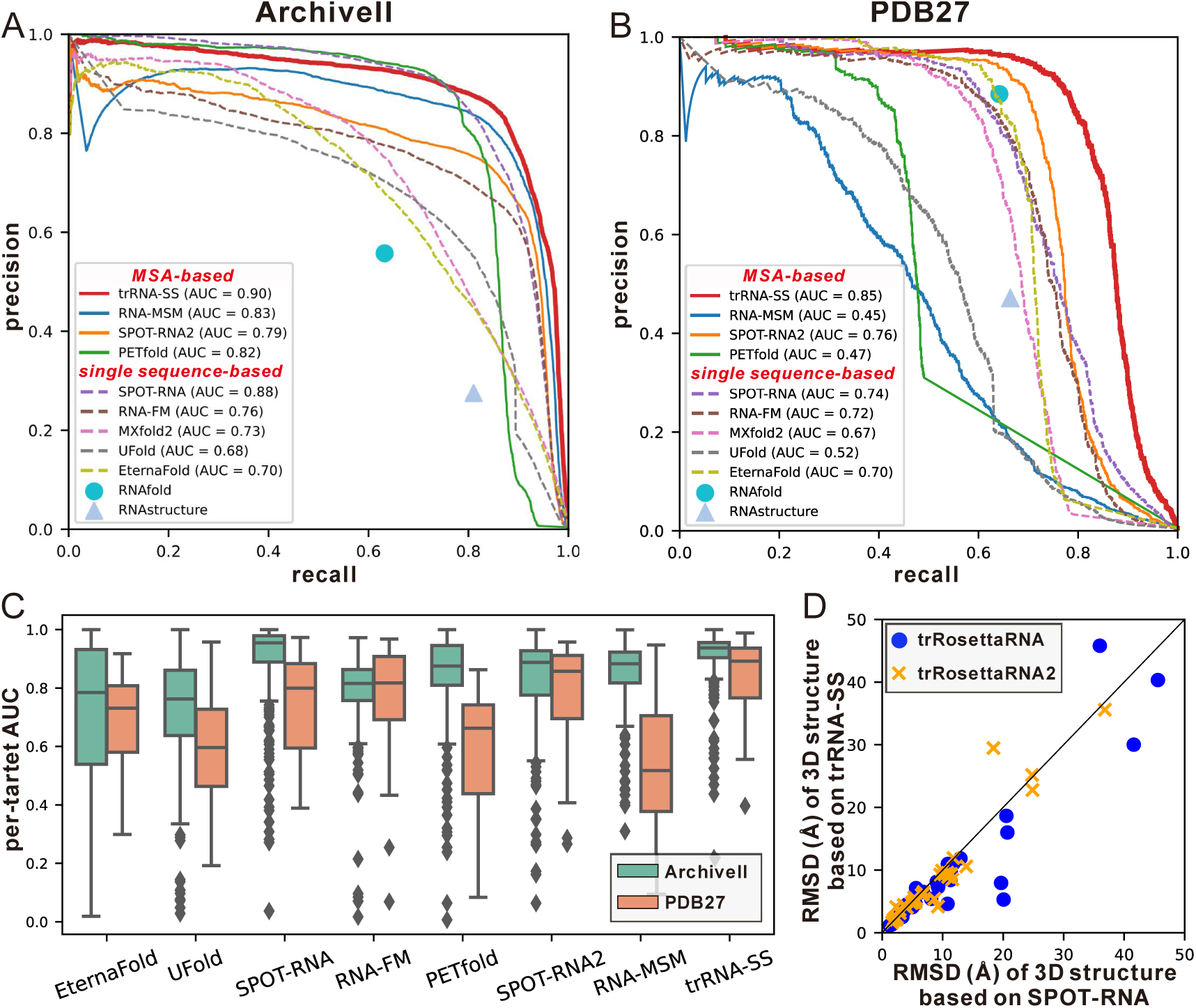
Performance of trRNA2-SS for RNA secondary structure prediction. (A, B) Precision-recall curves on ArchiveII and PDB27 test sets. (C) The distributions of per-target PR-AUC values. (D) Head-to-head comparisons between the 3D modeling results based on SPOT-RNA and trRNA2- SS.

Despite this challenge, trRNA2-SS maintains consistently robust performance across both datasets, achieving AUPRC values exceeding 0.8. Moreover, the relatively flat boxes associated with trRNA2-SS in Fig. 2C further illustrate the stability of its performance. These findings demonstrate the adaptability and robustness of trRNA2-SS, making it a state-of-the-art method for RNA secondary structure prediction.

We further examine whether the improved secondary structure prediction could enhance the 3D structure prediction. In our previous method trRosettaRNA, SPOT-RNA was the default method for providing input secondary structures. Fig. 2 has shown that trRNA2-SS can generate more accurate secondary structures compared to SPOT-RNA. Using these improved secondary structures as inputs, the average RMSD of trRosettaRNA on the TS39 set (the benchmark set for 3D structure prediction; see Methods) decreases by approximately 12% (from 10.1 Å to 8.9 Å). Notably, a significant majority of the RNAs (66.7%; 26 out of 39) exhibit improved 3D structure predictions (see Fig. 2D). This result confirms that the enhanced secondary structure prediction contributes to the improved accuracy of RNA 3D structure prediction.

### Application to RNAs without known secondary structures

The rapid inference powered by the deep learning model and GPUs enables the high-throughput prediction of secondary structures for a large volume of RNA sequences. As an application, we collect 98,008 non-redundant RNA sequences without known secondary structures from the RNAcentral database and predict their secondary structures with trRNA2-SS. The prediction accuracy was estimated using the confidence score C-ScoreSS defined in Eq. (8). A C-ScoreSS >0.75 strongly suggests an accurate prediction (Fig. S1). As shown in Fig. 3A, trRNA2-SS is more confident on the RNAs with well-defined functions, such as the regulatory *pre-miRNA*, the *signal recognition particle RNA* (SRP_RNA), and the *hammerhead ribozyme*, a well-known catalytic RNA. However, trRNA2-SS exhibits lower confidence for *long non-coding RNAs* (lncRNAs). This is consistent with the inherent characteristics of lncRNA, whose structure and function are largely influenced by interactions with other biomolecules, such as RNA, DNA, and proteins ^48^.

**Fig. 3.**
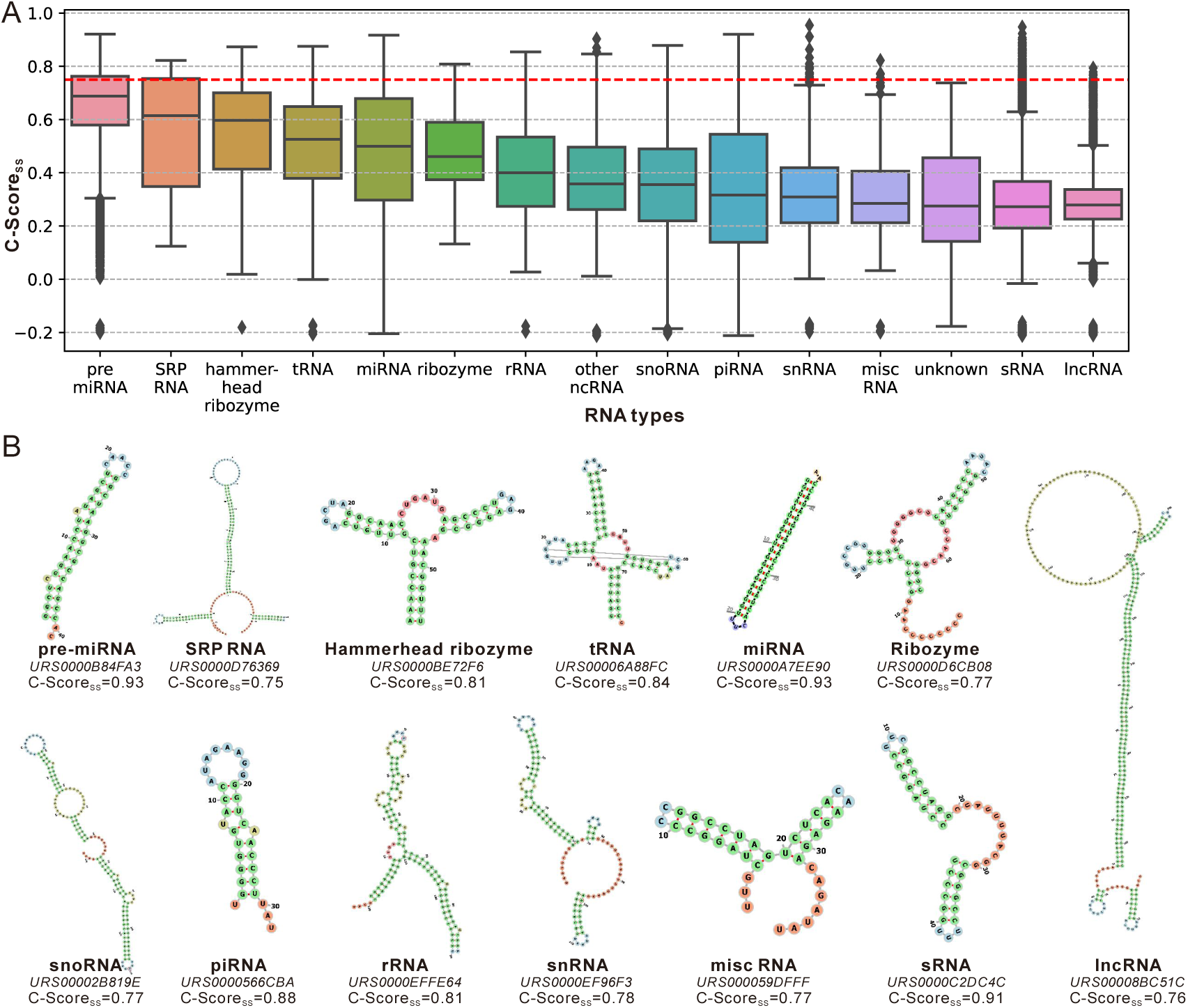
Application of trRNA2-SS to RNAcentral sequences without known secondary structures. (A) Distributions of the confidence score for predicted secondary structures (C-ScoreSS) on different RNA types. The red dashed line indicates the threshold (0.75) for a “confident” prediction. Each box encloses the middle 50% of the data (the Interquartile Range, IQR), with the bottom edge at the 25th percentile (Q1) and the top edge at the 75th percentile (Q3). The line inside the box indicates the median (50th percentile). The whiskers extend to the 1.5 * IQR from the box edges. Data points plotted individually outside the whiskers are outliers. (B) Representative examples of predicted secondary structures.

Despite this, trRNA2-SS can yield at least one confidence prediction for all related RNA types annotated by RNAcentral, including the challenging lncRNA. We present representative predictions for various RNA types in Fig. 3B. The corresponding predicted 3D structures are shown in Fig. S2. These predictions can serve as valuable structural references for related research. For example, for a human lncRNA (RNAcentral ID URS00008BC51C) located on chromosome 17, the trRNA2-SS prediction has a C-ScoreSS of 0.764, indicating a high-confidence prediction. According to its RNAcentral annotation, this RNA is an antisense lncRNA, implying that it relies on the interaction with other RNAs to perform biological functions, such as gene expression regulation and RNA processing. The secondary structure predicted by trRNA2-SS reveals a 70-nt loop, which may represent a potential binding site with other RNAs, thus providing valuable structural insight for investigating the function and mechanism of this lncRNA. To facilitate further research, we have made all confident trRNA2-SS predictions available on our website.

### Performance of trRosettaRNA2 on RNA 3D structure prediction

In addition to the SS prior module, another major advancement of trRosettaRNA2 over trRosettaRNA lies in its end-to-end (E2E) architecture powered by an SS-aware structure module. To assess the 3D structure prediction capabilities of trRosettaRNA2, we constructed a benchmark test set, TS39, which was carefully curated to ensure no redundancy with the training data (see Methods). For a comprehensive evaluation of performance, we assess multiple metrics, including global topology measures (RMSD and TM-score) and local accuracy metrics (lDDT and INF; see Methods for details).

To directly compare trRosettaRNA2 with the original trRosettaRNA, which takes SPOT-RNA-predicted secondary structures as input, we initially evaluate trRosettaRNA2 using secondary structure information from SPOT-RNA. This is achieved by bypassing the internal SS prior module and directly feeding the base-pairing probabilities predicted by SPOT-RNA into the RNAformer module. As shown in Fig. 4A and Table 1, trRosettaRNA2 (E2E) achieves an average RMSD of 8.2 Å on the TS39 dataset, a ∼18% improvement (lower RMSD) compared to trRosettaRNA. With the incorporation of the SS prior module, i.e., using the enhanced SS predictor trRNA2-SS, the average RMSD is further reduced to 7.8 Å. This demonstrates the effectiveness of the end-to-end architecture adopted in trRosettaRNA2.

**Fig. 4.**
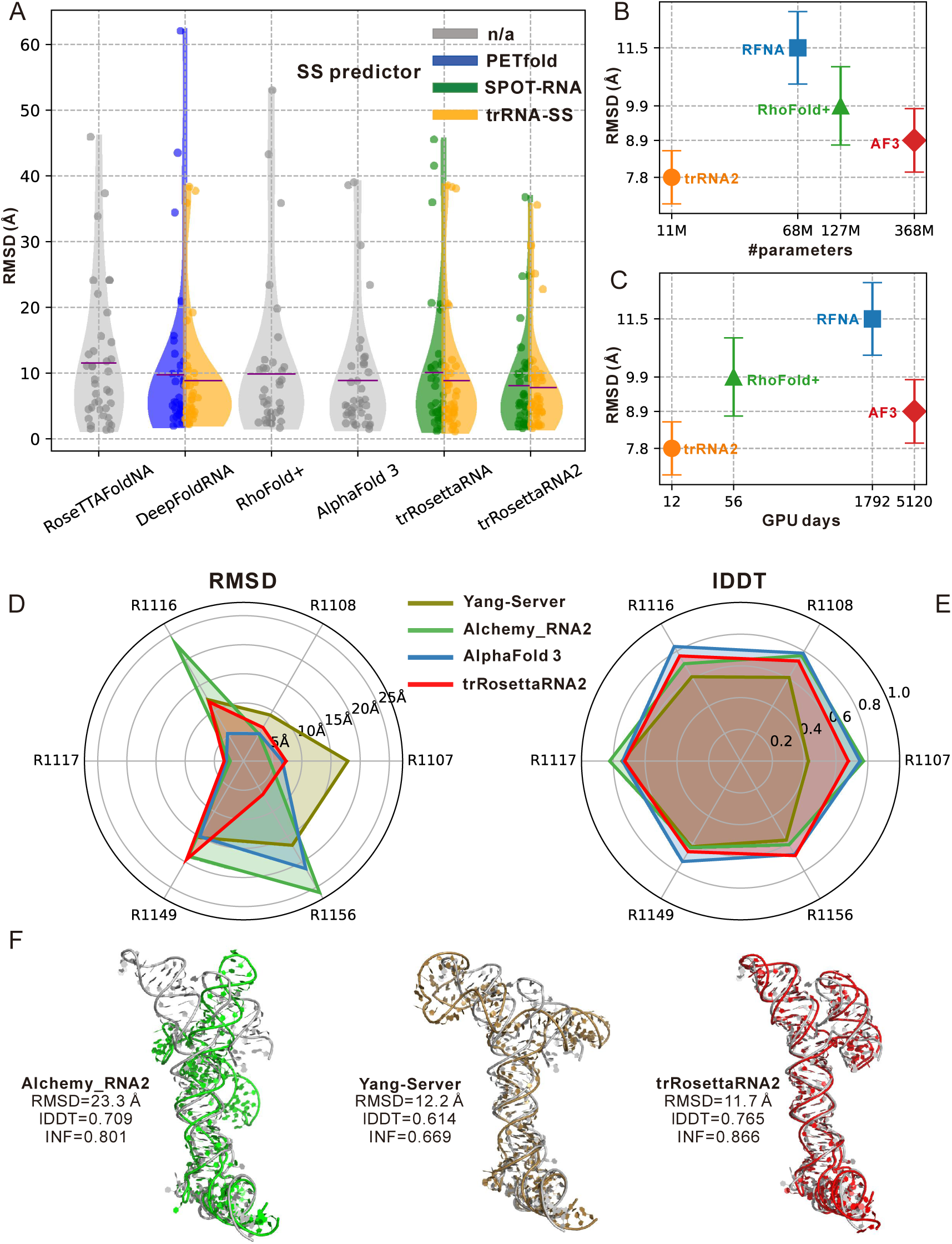
Performance of trRosettaRNA2 and other methods for RNA 3D structure prediction on TS39 dataset and CASP15 natural RNAs. (A) Comparison of RMSD distributions on TS39. Purple lines denote mean RMSD values. (B) Relationship between the average RMSD on TS39 and the number of parameters for end-to-end methods (trRNA2: trRosettaRNA2, RFNA: RoseTTAFoldNA, AF3: AlphaFold 3). (C) Relationship between the average RMSD on TS39 and training GPU days (i.e., GPU count × training days) for end-to-end methods. Note that the *x*-axes in panels (B) and (C) are logarithmically scaled for enhanced visualization. (D, E) Comparison of RMSDs and lDDTs on six natural RNAs. (F) Comparison of AIchemy_RNA2, Yang-Server, and trRosettaRNA on R1116. Note that we evaluate the “first” models for fairness.

**Table 1.** Comparison between trRosettaRNA2 and other representative methods on TS39 dataset. The best results are shown in bold, and the second-best results are underlined. ↓ = Lower is better; ↑ = Higher is better.

| Method | SS predictor | RMSD (Å) ↓ | IDDT ↑ | TM-score ↑ | INF ↑ | clashscore ↓ |
| --- | --- | --- | --- | --- | --- | --- |
| RoseTTAFoldNA | \ | 11.5 | 0.629 | 0.340 | 0.727 | 69.4 |
| DeepFoldRNA | PETfold | 9.7 | 0.678 | 0.341 | 0.745 | 18.1 |
|  | trRNA2-SS | 8.9 | 0.682 | 0.351 | 0.753 | 18.7 |
| RhoFold+ | \ | 9.9 | 0.654 | 0.321 | 0.711 | 38.2 |
| AlphaFold 3 | \ | 8.9 | 0.702 | **0.379** | **0.808** | 13.8 |
| trRosettaRNA | SPOT-RNA | 10.1 | 0.663 | 0.334 | 0.717 | 3.7 |
|  | trRNA2-SS | 8.9 | 0.688 | 0.348 | 0.722 | 3.0 |
| trRosettaRNA2<br>(E2E) | SPOT-RNA | 8.2 | 0.688 | 0.360 | 0.772 | 3.8 |
|  | trRNA2-SS | 8.0 | 0.699 | 0.362 | 0.775 | 4.4 |
|  | pLDDT top1 | <u>7.7</u> | 0.702 | 0.367 | <u>0.786</u> | 4.3 |
| trRosettaRNA2<br>(PyRosetta) | SPOT-RNA | 8.1 | 0.700 | 0.364 | 0.749 | 2.3 |
|  | trRNA2-SS | 7.8 | <u>0.711</u> | 0.364 | 0.759 | <u>2.0</u> |
|  | pLDDT top1 | **7.6** | **0.713** | <u>0.373</u> | 0.769 | **2.0** |

Interestingly, we observe that the PyRosetta version of trRosettaRNA2 is slightly more accurate than the E2E version, likely due to the enhanced structural plausibility achieved through energy minimization. Considering that the energy minimization procedure is primarily guided by the 2D geometries predicted by the neural network, this result suggests that the end-to-end training also boosts the 2D geometries prediction. In practice, for regular RNAs, the E2E version provides a fast way to generate satisfactory structures. However, for large RNAs (>400 nucleotides), where the neural network may face challenges in resolving structural violations, the PyRosetta version can yield more structurally plausible results.

In trRosettaRNA2, the confidence score of the predicted 3D structure is directly generated by the neural network, in the form of a predicted lDDT (pLDDT) score. As illustrated in Fig. S3A, the pLDDT score exhibits a strong correlation with the real RMSD values, achieving a Pearson correlation coefficient (PCC) of -0.773. To demonstrate its practical utility, we employ the pLDDT score to select optimal models between trRosettaRNA2 predictions based on SPOT-RNA and trRNA2-SS. The results reveal that this pLDDT-guided selection strategy effectively enhances prediction accuracy for both the E2E version and the PyRosetta version (Table 1 and Fig. S3B). These findings highlight the potential for further improvements in trRosettaRNA2 performance through the systematic exploration of diverse inputs.

We also observed that the recycling mechanism can help improve performance especially when the input secondary structure is inaccurate. As shown in Fig. S4, while recycling enhances the INF of predicted 3D structures across most input SS AUC levels, the most pronounced improvements occur at the lowest AUC level (<0.4). This demonstrates the efficacy of the recycling mechanism in compensating for prior base-pairing information. Furthermore, incorporating SS information into the structure module (i.e., the SS-aware structure module) also boosts performance for targets with low SS AUC, revealing that even incomplete base-pairing information can help the structure module yield improved results for challenging targets.

To illustrate the effect of the recycling mechanism more specifically, we examine an example RNA with PDB ID of 8YDC, a pseudoknot-containing *hammerhead ribozyme*. As shown in Fig. S5A, SPOT-RNA failed to identify several interactions crucial for the 3D topology of this RNA (e.g., the triple helix formed between residues 3∼7, 14∼18, and 19∼23, highlighted in dashed circle). Consequently, trRosettaRNA, using this limited SS input, could not accurately model these interactions (Fig. S5B), resulting in a 3D structure with high RMSD (>20 Å; Fig. S5C). In contrast, while trRosettaRNA2 initially struggled to capture SPOT-RNA-missed interactions within a single cycle, the final recycled prediction (cycle=4) successfully resolved this, resulting in a substantial RMSD reduction (Fig. S6, RMSD dropping from ∼15 Å to ∼6 Å). Notably, the complete trRosettaRNA2 model, using trRNA2-SS input instead of SPOT-RNA, achieves acceptable accuracy even without recycling (∼8 Å), while recycling provides only an incremental improvement (∼2 Å). This observation illustrates that while enhanced prior knowledge is beneficial, trRosettaRNA2’s recycling mechanism ensures model robustness when the base-pairing information from the input SS is limited or inaccurate.

### Comparison of trRosettaRNA2 and other methods

In our previous work trRosettaRNA, we have demonstrated that deep learning-based methods outperform traditional automated methods relying on physical energy and/or fragment assembly. Thus, in this study, we only compare trRosettaRNA2 with other state-of-the-art deep learning-based methods, including DeepFoldRNA and RoseTTAFoldNA (published around the same time as trRosettaRNA), as well as more recently published methods RhoFold+ and AlphaFold 3 (AF3). Unless otherwise specified, all methods were run locally using identical inputs (MSA and/or SS).

As shown in Table 1, trRosettaRNA2 exhibits competitive performance with AF3 and outperforms other methods across both metrics. Specifically, trRosettaRNA2 achieves better RMSD and lDDT than AF3, though with slightly lower TM-score and INF, while still outperforming all other representative methods across all metrics. Furthermore, trRosettaRNA2 models demonstrate fewer steric clashes, highlighting the effectiveness of its fine-tuning strategy and energy-based relaxation/minimization.

One of the key advantages of trRosettaRNA2 lies in its parameter efficiency. As shown in Fig. 4B, trRosettaRNA2 achieves competitive performance with only ∼11M parameters. This represents a significant reduction compared to other end-to-end methods: only 1/6 of RoseTTAFoldNA, 1/11 of RhoFold+, and 1/33 of AF3. This efficiency is also observed in training cost, as the training of trRosettaRNA2 required only ∼12 days on a single A100/A800 GPU, much fewer than RhoFold+ (∼1 week on 8 A100 80GB GPUs), RoseTTAFoldNA (∼4 weeks on 64 GPUs), and AF3 (∼20 days on 256 A100 GPUs). These findings highlight the remarkable efficiency of trRosettaRNA2.

During the preparation of our manuscript, a new deep learning-based method, DRfold2 ^49^, was proposed. For each target RNA, DRfold2 employs a composite language model to embed the nucleotide sequence and uses 80 Transformer-based neural network models to predict 80 sets of backbone frames and 2D geometries. These predictions are subsequently used to generate 3D structures through the CSOR protocol (Clustering, Scoring, Optimization, and Refinement). To compare trRosettaRNA2 with DRfold2, we further filtered the TS39 set to exclude sequences redundant to the DRfold2 training set (based on an E-value threshold of 10 by BLASTN ^50^ search), resulting in a subset of 34 RNAs.

As shown in Table S1, trRosettaRNA2 achieves a lower average RMSD than DRfold2 on these 34 RNAs (10.2 Å vs. 10.7 Å) and remains competitive across other accuracy metrics. Notably, the steric clashes in trRosettaRNA2 predictions are significantly fewer than those in DRfold2 predictions, with the average clashscores of 2.2 and 35.3, respectively. Note that the default DRfold2 prediction is based on an ensemble of 80 neural network models, whereas trRosettaRNA2 utilizes only a single model. Considering this difference in approach, we evaluate the best-performing single model of DRfold2 on these 34 RNAs, which demonstrates inferior performance compared to trRosettaRNA2 in terms of both RMSD and lDDT. Overall, trRosettaRNA2 exhibits a more favorable balance between modeling accuracy and structural plausibility compared to DRfold2.

Furthermore, the design of the SS prior module allows trRosettaRNA2 to enhance performance by exploring diverse prior information (e.g., the combination of the SS from SPOT-RNA and trRNA2-SS further improves accuracy), whereas AF3 and DRfold2 rely solely on the co- evolutionary signal extracted by MSA or language model. This flexibility also helped our group (Yang-Server) achieve superior results in the CASP16 competition (discussed later).

### Performance on CASP15 RNA targets

In 2022, we participated in the CASP15 RNA structure prediction experiments using the automated method trRosettaRNA. This blind assessment demonstrated that trRosettaRNA (Yang-Server) achieved competitive RMSD compared to the top human groups (e.g., AIchemy_RNA ^51^) for natural RNAs. However, its performance on local metrics, such as lDDT and INF, was less competitive, revealing inaccuracies in the local structural details of trRosettaRNA models. Though the Yang-Server models for these targets achieved lower RMSD than the AIchemy_RNA2 models, their local metrics (lDDT and INF) lagged significantly behind those of AIchemy_RNA2.

In this study, we benchmark trRosettaRNA2 on six natural RNAs from CASP15 (excluding R1189 and R1190 due to their complex interactions with proteins that could skew the evaluation).

As shown in Table S2 and Figure 4D, trRosettaRNA2 consistently outperforms Yang-Server (powered by trRosettaRNA) and demonstrates competitive performance with AF3, aligning well with the results observed on the TS39 dataset. Notably, trRosettaRNA2 also surpasses the leading human group, AIchemy_RNA2, in terms of RMSD, while maintaining competitive performance across other metrics, which also represents an enhancement compared to trRosettaRNA. We highlight this enhancement using a challenging target R1116, an *Enterovirus cloverleaf RNA* (PDB ID: 8S95). Figure 4E reveals that the AIchemy_RNA2 model exhibits inaccuracies in the cloverleaf region and its orientation relative to the tRNA component, with an RMSD exceeding 20 Å. The Yang-Server model displays improved global topology (RMSD of 12 Å), but its local interaction details are inferior to AIchemy_RNA2, as indicated by lower lDDT and INF values. In contrast, the trRosettaRNA2 model demonstrates better accuracy than AIchemy_RNA2 in both global topology and local details, achieving a 49.8% lower RMSD, a 7.9% higher lDDT, and an 8.1% higher INF compared to AIchemy_RNA2. These findings further confirm the robust enhancements of trRosettaRNA2 over its predecessor, trRosettaRNA.

Automated modeling, however, continues to face challenges with the four synthetic RNAs, which are characterized by intricate topologies and a scarcity of homologous sequences. As shown in Table S3, all evaluated automated methods, including AF3, consistently exhibit suboptimal performance on these RNAs. While trRosettaRNA2 demonstrated a relatively better ability to approximate global topology as an automated approach (indicated by its lower RMSD and higher TM-score), its overall accuracy remained inadequate and significantly lagged behind the human expert performance. This persistent performance gap illustrates the inherent limitations of current automated approaches when confronted with highly complex and novel RNA structures, highlighting the necessity for future advancements in the collaboration of computational methodologies and wet-lab experiments to effectively tackle these challenging edge cases.

### Blind test results in CASP16

Based on trRosettaRNA2, we participated in the blind tests of the CASP16 RNA prediction experiment as two automated server groups, Yang-Server (TS052) and Yang-Multimer (TS456). Yang-Server was optimized for submitting the best predictions using diverse SS resources, which are trivial to obtain; while Yang-Multimer used the default SS predictions from trRNA2-SS to compare with the other four deep learning predictors.

### Yang-Server results

According to the official ranking for the 36 RNA monomer targets (https://predictioncenter.org/casp16/zscores_rna.cgi), Yang-Server is the top-performing server group (1/16), either ranked by the “first” models or the “best” models, and is surpassed only by three human groups (i.e., 4-th among 64 participating groups). In contrast, AF3-server (powered by AF3; TS304) ranks 9-th among the 64 groups.

As mentioned in our previous work ^13^, trRosettaRNA’s performance can be enhanced by exploring template secondary structures as input restraints for three RNAs from CASP15. Building upon this finding, we prepared multiple input SSs for each target during CASP16. These SSs included predictions from various algorithms (trRNA2-SS, SPOT-RNA, and EternaFold) as well as those derived from available templates (detected by RNAthreader ^52^ and R2DT ^53^). Additionally, we trained a variant of trRosettaRNA2 that does not use the SS prior module, providing an SS-free prediction. For several targets (R1221s2, R1224s2, R1281, R1289, R1291) with identifiable 3D templates (using RNAthreader), we also performed template-based modeling to expand the decoy set. Finally, for each target, we used an in-house RNA quality assessment (QA) method (Liu et al, in preparation) to select the top 5 models from the decoy set for submission as Yang-Server (TS052).

Considering that trRosettaRNA2 was trained with a maximum crop size of 384 nucleotides due to the limited computational resources, its applicability to larger RNAs (>400 nucleotides) may be hindered. Therefore, for the large RNA targets without available templates in CASP16 (R1241, R1248, R1250-R1254, R1283, R1285, R1286, R1290), we chose to submit the AF3 predictions, as its training utilized a larger crop size of 768. For the remaining 23 targets, Yang-Server still ranks as the best server group (see Fig. 5A). As shown in Table S4, the “best” models for these 23 targets from Yang-Server outperform those from AF3-server across global metrics (RMSD, TM-score, and GDT-TS) and demonstrates competitive performance in local metrics (lDDT and INF), while achieving a much lower clashscore.

**Fig. 5.**
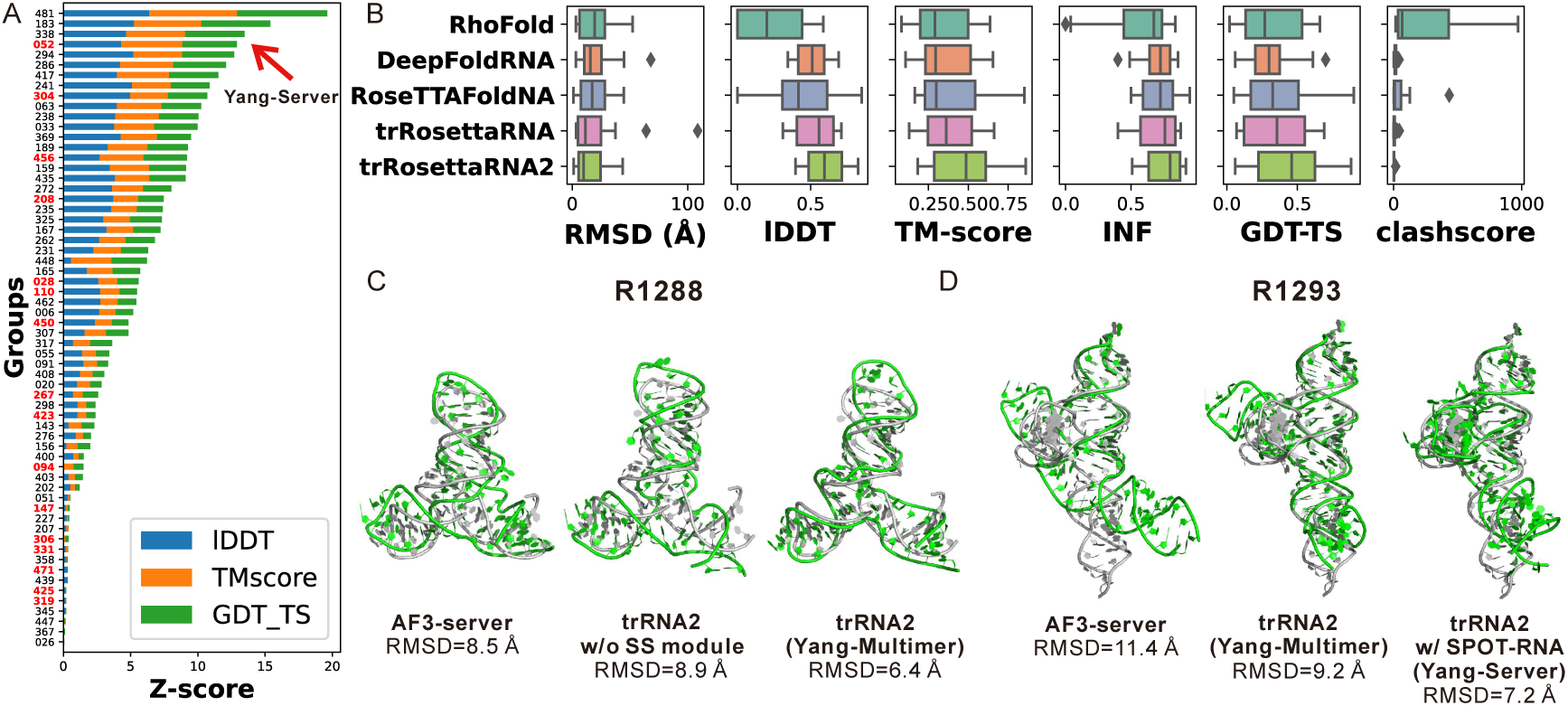
CASP16 blind test results. (A) Ranking of the summed Z-score (>0; calculated according to CASP16 official guidelines) for the 23 RNA targets shorter than 400 nucleotides or with templates. Automated server groups are highlighted in bold red on the *y*-axis tick labels. (B) Results for blind benchmark test performed by Yang-Multimer submission (note that the RhoFold and RoseTTAFoldNA predictions were not relaxed). (C, D) Two examples to illustrate the comparison between trRosettaRNA2 (trRNA2) and AlphaFold 3 (AF3-server). Predicted structures (in green) are superimposed onto the experimental structures (in gray).

### Yang-Multimer results

To conduct a more direct blind benchmark test of trRosettaRNA2, we simultaneously submitted multiple predictions through another server group, Yang-Multimer (TS456). Specifically, we submitted the predictions from trRosettaRNA2, trRosettaRNA, RoseTTAFoldNA, DeepFoldRNA, and RhoFold as models 1∼5, respectively, within the Yang-Multimer group. All five methods used the same inputs (MSA and/or SS). According to the CASP16 official rankings, Yang-Multimer places 5-th among the 16 server groups ranked by the summed Z-score (>0) of the “first” models. Note that only 23 targets with Yang-Multimer submissions are included in the official ranking, which means that the Z-scores for the remaining 13 targets are both zero. If ranked by the average Z-score, Yang-Multimer is the second-best server group (behind Yang-Server) among the 37 groups (10 server groups) with more than 20 submissions, reflecting the superior performance of trRosettaRNA2.

Fig. 5B and the top portion of Table 2 summarize the results of Yang-Multimer. Among the Yang-Multimer submissions, trRosettaRNA2 outperforms the other four methods across all metrics. Furthermore, we assess the recently published RhoFold+ ^16^ on a subset of 20 targets for which experimental structures are available. The results indicate that RhoFold+ performs closely to RhoFold, while trRosettaRNA2 outperforms RhoFold+ on all metrics. These observations are consistent with the benchmark results on the TS39 dataset discussed earlier. In addition, the trRosettaRNA2 predictions displayed significantly fewer steric clashes compared to other methods, further highlighting its advantage in both model accuracy and physicochemical plausibility.

**Table 2.** Performance on CASP16 RNA monomer targets with Yang-Multimer submissions. All metrics are sourced from the official CASP16 evaluation unless specified otherwise.

| Group | Method | RMSD<br>(Å) ↓ | IDDT<br>↑ | TM-score<br>↑ | GDT-TS ↑ | INF<br>↑ | clashscore<br>↓ |
| --- | --- | --- | --- | --- | --- | --- | --- |
| Yang-Multimer | RhoFold <sup>#</sup> | 19.9 | 0.224 | 0.328 | 0.298 | 0.562 | 223.9 |
|  | DeepFoldRNA | 20.1 | 0.509 | 0.345 | 0.312 | 0.701 | 14.8 |
|  | RoseTTAFoldNA<sup>#</sup> | <u>18.8</u> | 0.428 | <u>0.387</u> | <u>0.355</u> | <u>0.718</u> | 44.4 |
|  | trRosettaRNA | 20.1 | <u>0.530</u> | 0.385 | 0.351 | 0.708 | <u>6.9</u> |
|  | trRosettaRNA2 | **15.9** | **0.598** | **0.464** | **0.432** | **0.759** | **3.2** |
| Yang-Server | First model | 13.8 | 0.635 | 0.510 | 0.464 | 0.790 | <u>3.7</u> |
|  | Best model | **10.2** | <u>0.663</u> | **0.548** | **0.496** | <u>0.815</u> | **1.4** |
| AF3-server | First model | 12.4 | 0.659 | 0.504 | 0.457 | 0.809 | 16.3 |
|  | Best model | <u>11.3</u> | **0.675** | <u>0.527</u> | 0.478 | **0.818** | 11.5 |
| Reduced set * | RhoFold <sup>#</sup> | 19.6 | 0.496 | 0.317 | 0.313 | 0.522 | 199.6 |
|  | RhoFold+ | 25.6 | 0.512 | 0.317 | 0.320 | 0.636 | 65.7 |
|  | trRosettaRNA2 | 14.3 | 0.636 | 0.443 | 0.429 | 0.758 | 2.1 |

\* 20 targets with available experimental structures. The metric values on this set are slightly different from the official results as we manually calculated them.
<sup>#</sup> RhoFold and RoseTTAFoldNA predictions were not relaxed.

### Case study for CASP16

As illustrated above, trRosettaRNA2 achieves competitive results compared to AF3, despite requiring significantly fewer computational resources for training. This can be attributed to the incorporation of prior information through the SS prior module in trRosettaRNA2, leveraging the fact that SSs are generally more readily predictable than complex 3D structures. Figs. 5C and 5D present two examples to illustrate this.

The first example is R1288, a stabilized *S-adenosylmethionine analogue-utilizing ribozyme* (SAMURI; Fig. 5C) which bound to SAH and two Mg²⁺ ions. For this target, trRosettaRNA2 achieved similar accuracy with the best model from AF3-server when no SS prior information was used (8.9 Å and 8.5 Å, respectively). With the SS prior module incorporated, the RMSD of the trRosettaRNA2 model was improved to 6.4 Å. This example illustrates the advantage of leveraging SS prior information, leading to improved results compared to the methods that do not support this, such as AF3.

The second example is R1293, which comes from a *translation enhancer motif* M1293, a complex composed of two protein subunits and one RNA subunit. For this target, while the default trRosettaRNA2 model outperforms AF3 (i.e., Yang-Multimer vs. AF3-server), the performance can be further enhanced when the SPOT-RNA SS is used as input, resulting in an even more accurate model for Yang-Server compared to Yang-Multimer. This example highlights the potential of trRosettaRNA2 to explore diverse SS prompts to yield more accurate predictions.

Note that the above two RNA targets are both involved in complexes with other molecules: R1288 is bound to ligands, and R1293 interacts with proteins. The prediction of these entire complex structures falls outside the scope of our method, which is a limitation of our approach compared to AF3. However, we observe that incorporating binding partner information does not improve the performance of AF3 for these two RNA subunits (Fig. S7A). For instance, despite AF3 predicting a nearly correct binding site for R1293, the overall bound conformation exhibits significant deviations from the experimental structure, with a notable mis-twisting of the junction region (Fig. S7B). This reflects that accurate prediction of intermolecular interactions, particularly those involving RNA, remains a challenging problem, possibly due to a scarcity of relevant training data.

### Exploring the conformational landscape of RNase P RNA

Compared to protein, RNA structures are typically more flexible and dynamic, often existing as a heterogeneous conformational ensemble rather than a single static structure ^54–56^. However, modeling this inherent dynamic nature remains a challenge for current computational methods. Building up our finding that SS inputs modulate trRosettaRNA2’s output, we apply trRosettaRNA2 to predict the heterogeneous structures of the ribonuclease P (RNase P) RNA. This RNA consists of a highly-conserved catalytic domain (C-domain) and a specificity domain (S-domain) known for significant structural heterogeneity across different species. Elucidating the S-domain’s structural variability is crucial for understanding the catalytic activity and mechanism of RNase P. Recently, 158 distinct RNase P RNA conformations were determined via deep learning analysis of atomic force microscopy (AFM) images ^26^. This experimentally observed ensemble shows significant diversity, with an average root mean square fluctuation (RMSF) reaching 19.1 Å (Fig. 6A). The variability is significantly more pronounced in the S-domain (average RMSF 31.1 Å) than in the C-domain (average RMSF 11.7 Å), a trend clearly evident in the per-residue RMSF plot (Fig. 6B).

**Fig. 6.**
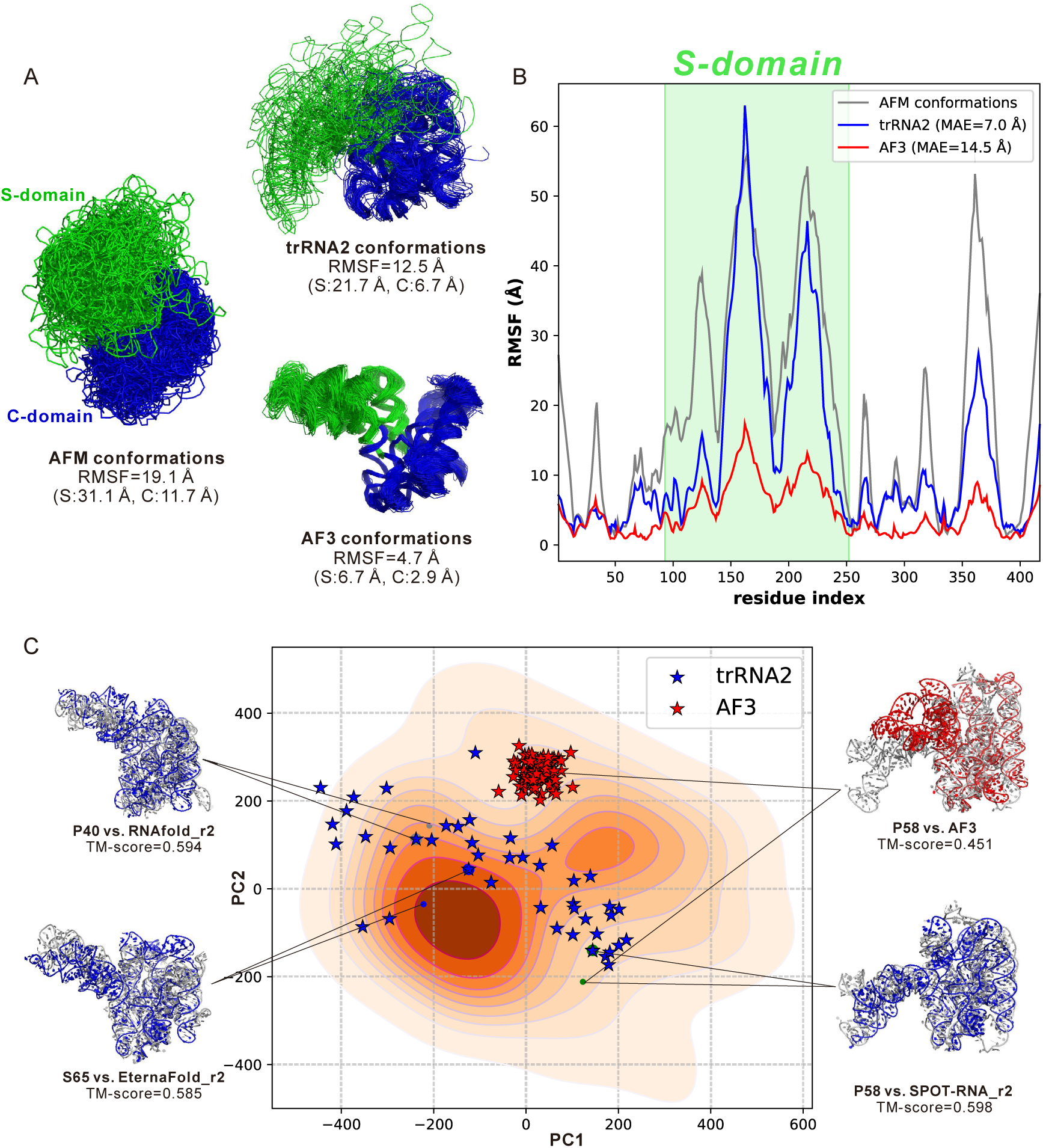
Exploring the RNase P RNA conformational landscape. (A) Comparison of conformation ensembles determined by AFM versus those predicted by trRosettaRNA2 (trRNA2) and AF3. The conformations have been superimposed onto a reference structure (PDB: 2A64). Parentheses show average S/C-domain RMSFs. (B) Per-residue RMSF plot, with the S-domain shaded green. (C) Visualization of the conformational landscape via Principal Component Analysis (PCA) based on C1’ atom positions. For each highlighted example, predicted structures (trRNA2: blue; AF3: red) are superimposed onto the corresponding experimental conformation (gray). The trRNA2 predictions are named using the format “{SS predictor}_r{recycling times}”.

To sample heterogeneous conformations using trRosettaRNA2, we generate predictions using 11 distinct SSs as input. These SSs were obtained from various tools: trRNA2-SS (along with its three single-sequence models), SPOT-RNA, SPOT-RNA2, EternaFold, MXfold2, RNAfold, RNAstructure, and a template SS detected by R2DT. For each SS input, four structures were predicted using the E2E version of trRosettaRNA2 (corresponding to its four internal cycles), while the input MSA was kept fixed. As a comparison, we also explore AF3’s ability to capture conformational heterogeneity by running AF3 with 20 different random seeds (5 models per seed). In total, we generate 44 trRosettaRNA2 predictions and 100 AF3 models for RNase P RNA.

trRosettaRNA2 demonstrates a much better ability to capture structural heterogeneity. As shown in Fig. 6A, the trRosettaRNA2 predictions exhibit significant dynamics (overall RMSF=12.5 Å), particularly within the S-domain (RMSF>20 Å). Furthermore, its per-residue RMSF profile aligns well with that derived from the experimental conformations (Fig. 6B), achieving a PCC of 0.894. Notably, trRosettaRNA2 accurately captures the high flexibility observed in the two most dynamic stem loops within the S-domain (residues ∼150-179 and ∼205-235). In contrast, the AF3 predictions are highly homogeneous across both domains, with an overall RMSF of only ∼5 Å. Compared to the experimental conformations, AF3 significantly underestimates the conformational fluctuation for almost all residues (Fig. 6B). These results highlight the advantage of trRosettaRNA2 over AF3 for predicting conformational ensembles.

The potential of trRosettaRNA2 is further underscored by the conformational landscape visualization in Fig. 6C. Despite generating twice as many structures, AF3’s predictions are highly concentrated within a small area (red stars), whereas trRosettaRNA2’s predictions exhibit greater diversity and spread. For instance, for the three structurally distinct conformations highlighted in Fig. 6C, trRosettaRNA2 successfully generates accurate predictions via different SS inputs. This result likely stems from trRosettaRNA2’s ability to leverage diverse SS inputs, a mechanism not supported by AF3. As a specific example, for the P58 conformation, the closest AF3 prediction remains distant in the landscape, yielding a TM-score of only 0.451, considerably lower than the trRosettaRNA2 prediction (∼0.6). These findings illustrate the potential of trRosettaRNA2 for applications in studying RNA conformational landscapes, a crucial and challenging area in RNA biology.

### Ablation study

In this section, we analyze the contribution of the SS prior module and MSA embedding to the performance of trRosettaRNA2. To achieve this, we perform an ablation study by individually removing each module and retraining the neural network while keeping all other configurations identical. Additionally, we also assess the impact of the two fine-tuning stages (violation fine-tuning and resampled fine-tuning; see Methods), utilizing checkpoints saved after each training phase. During inference, all ablation models are provided with the same input MSAs (except for the “w/o MSA embedding” model, which uses the single sequence only). For simplicity, 3D structures were generated using the end-to-end (E2E) approach rather than the PyRosetta approach.

### Impact of SS prior module

As shown in Fig. 7A, the most significant performance decline is observed when the SS prior module is removed. The average RMSD on the TS39 set increases from 8.0 Å to 10.2 Å. Notably, for 76.9 % (30/39) of RNAs in TS39, the removal of the SS prior module leads to higher RMSD values. All other accuracy metrics, including lDDT, TM-score, and INF, also exhibit significant deterioration. This indicates the importance of the base-pairing prior knowledge embedded within the extensive secondary structure data.

**Fig. 7.**
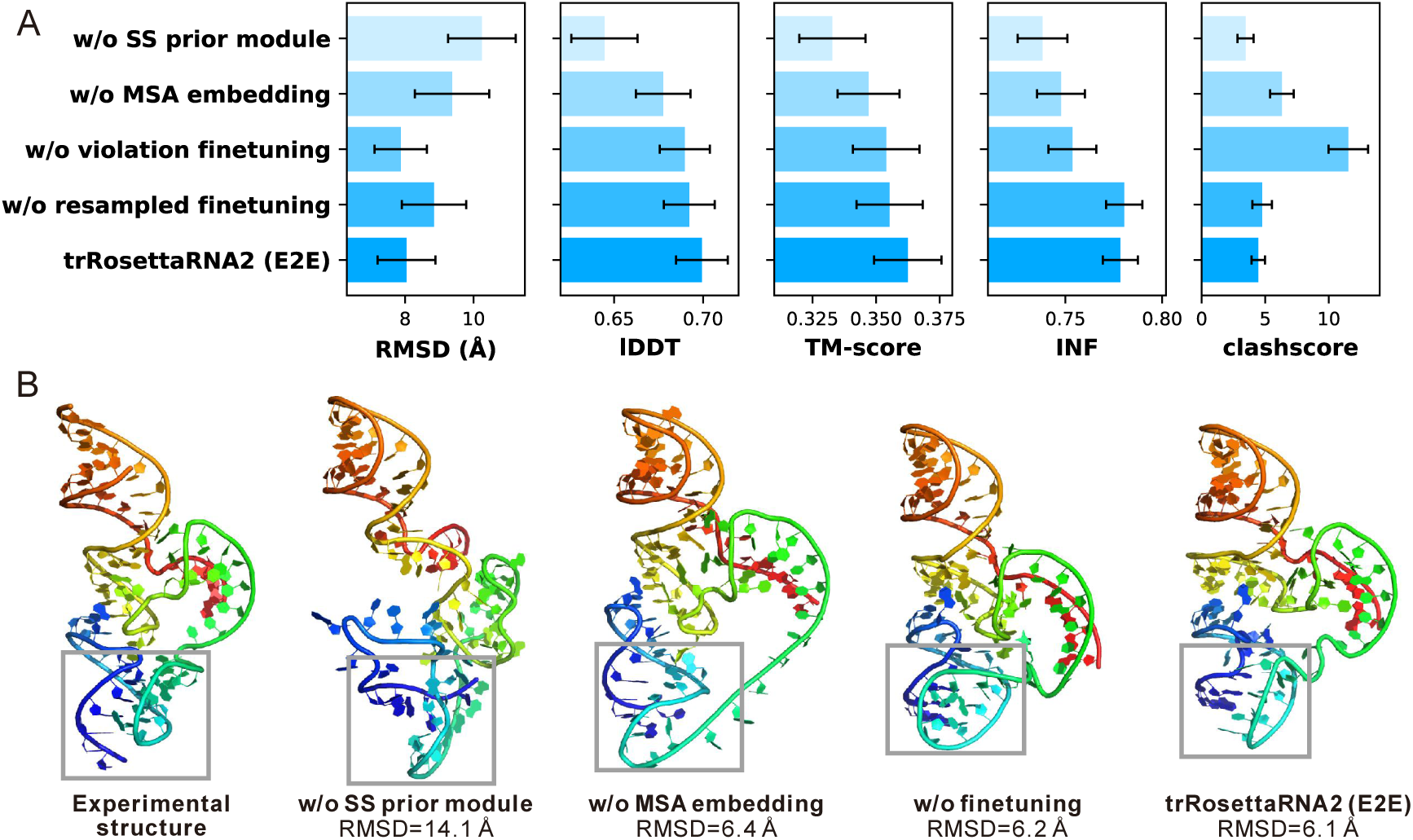
Ablation study of trRosettaRNA2. (A) Comparison of ablation models on the TS39 dataset. Each bar represents the mean value of the corresponding metrics across the TS39 set. Each error bar indicates one-tenth of the standard deviations. (B) 3D modeling results for the example RNA (PDB ID: 8YDC). The gray boxes highlight the triple helix which is crucial for the 3D topology of this RNA.

Further investigation reveals weak correlations between the performance degradation and the quality of SSs generated by the SS prior module (PCC -0.26∼0.003; Fig. S8). Notably, even for the targets with low-quality SSs, the removal of the SS prior module still worsens the 3D modeling accuracy. This finding is consistent with our earlier analysis of the SS-aware structure module, highlighting the value of extra prior information for modeling challenging targets.

### Impact of MSA embedding

Removing the MSA embedding also exhibits a detrimental effect, although less pronounced than that of excluding the SS prior module. The average RMSD increases from 8.0 Å to 9.4 Å. For 76% of the 25 RNAs (19/25) possessing more than one effective homologous sequence (i.e., *Neff*>1), the removal of MSA embedding worsens modeling accuracy. For the remaining 14 RNAs lacking effective homologous sequences (i.e., *Neff* =1), the impact was less consistent: the performance degradation is observed in only half of the RNAs, with the other half exhibiting minimal improvements. This result illustrates the importance of the co-evolutionary signals implied in the MSA.

However, according to Fig. S8, the correlations between the accuracy degradation and the MSA quality (measured by the logarithm of *Neff*) are also weak (PCC -0.08∼0.2). Considering the overall average RMSD, the introduction of MSA embedding proves beneficial for both the *Neff* = 1 and *Neff* > 1 groups. This finding suggests that MSA embedding may also contribute to training efficiency, potentially enhancing inference performance even when the input MSA quality is limited.

### Impact of violation fine-tuning

The first fine-tuning stage of trRosettaRNA2 aims to minimize steric clashes and bond violations in the predicted structures. As shown in Fig. 7A, this stage significantly reduces clashes, with the average clashscore decreasing from 11.5 to 4.4, confirming the effectiveness of this process. The reduction of clash is more obvious for targets that initially exhibit higher clashscores (PCC = 0.944; Fig. S8). Concurrently, the average RMSD remains largely unchanged, while both the lDDT and TM-score show slight improvements.

Remarkably, the improvement in the INF metric also shows a significant correlation with the initial clashscore (PCC=0.557). As depicted in Fig. S8, the violation fine-tuning process results in improved interaction networks for all RNAs with an initial clashscore > 20, and notably, two of these RNAs achieve an INF metric improvement of over 0.2. This may be attributed to the more realistic local structures generated by the fine-tuned model.

### Impact of resampled fine-tuning

The final training stage of trRosettaRNA2 is designed to enhance performance on challenging RNAs by increasing the sampling weights for RNAs with low lDDT scores. According to Fig. 7A, this fine-tuning stage results in improvements in RMSD, lDDT, and TM-score, while the INF and clashscore remain relatively stable. Among the 39 RNAs, 17 exhibited an RMSD > 6 Å before the resampled fine-tuning stage. After this stage, a majority (82.4%, or 14 out of 17) of these “hard” targets achieved an improvement in RMSD, a notably higher percentage compared to the remaining “easy” targets (59.9%, or 13 out of 22). This result confirms the effectiveness of our resampled fine-tuning strategy for improving the prediction accuracy of difficult RNA targets.

### Case study for ablation

We reuse the example RNA (PDB ID:8YDC) to specifically illustrate the contribution of each component (Fig. 7B). Apart from the pseudoknot, the triple helix formed by residues 3∼7, 14∼18, and 19∼23 (indicated by gray boxes in Fig. 7B) is also crucial for the overall 3D topology.

Without the SS prior module, the model successfully predicted the helix near the 5’ terminus. However, the pseudoknot and the triple helix were not built well, resulting in an inaccurate 3D structure with an RMSD of 14.1 Å. For the “w/o MSA embedding” model, the pseudoknot was captured well, and the 3D structure was significantly more accurate with an RMSD of 6.4 Å, perhaps due to the contribution of the SS prior module. Nevertheless, the triple helix region remained incomplete, leading to an unnatural straight loop of 5 nucleotides. When both the SS prior module and MSA embedding were included, the model generated a more reasonable 3D structure that captured all key interactions, although the orientation of the triple helix was still incorrect. Finally, with the two-stage fine-tuning, the complete trRosettaRNA2 model predicts a more accurate 3D structure, achieving a lower RMSD of 6.1 Å. This example further demonstrates the effectiveness of the network architecture and training strategy employed by trRosettaRNA2.

## DISCUSSION

We develop trRosettaRNA2, an automated end-to-end algorithm for RNA 3D structure prediction. Rigorous benchmark testing on two test sets, along with blind assessments during CASP16, demonstrates that trRosettaRNA2 outperforms its predecessor, trRosettaRNA, and other representative deep learning-based methods. Notably, trRosettaRNA2 achieves comparable performance to AlphaFold 3, despite utilizing considerably fewer parameters and computational resources for its development and training. This enables our group, Yang-Server, to achieve the top ranking among automated server groups in CASP16.

According to the ablation study, the most significant contribution stems from the secondary structure module. This module provides crucial prior information about base-pairing, which helps mitigate performance degradation caused by the scarcity of 3D structure data. Furthermore, this module can also be used as an independent secondary structure predictor, which we have named trRNA2-SS. Benchmark tests have shown that trRNA2-SS achieves state-of-the-art performance in secondary structure prediction. As a practical application, trRNA2-SS is used to predict the secondary structure for nearly 100,000 RNAcentral sequences lacking known SS, yielding high-confidence predictions across various RNA types. These high-confidence predictions can serve as a valuable structural reference for related research.

However, the automated prediction of the four synthetic RNAs from CASP15 remains challenging. This may be due to the complicated and novel interaction architectures within these RNAs. Despite this, our CASP16 report (Wang et al, 2025) indicates that incorporating template information helped our group, Yang-Server, to achieve the most accurate prediction for R1281, a mutant of the most intricate synthetic RNA in CASP15 (R1138). This result highlights the potential of integrating template information with deep learning-based methods to tackle challenging RNA targets. Moreover, the combination of deep learning and advanced experimental techniques, such as cryo-EM and atomic force microscopy (AFM), has shown great promise in determining RNA structures and the RNA conformational space ^25, 57^. Considering the inherent difficulties in modeling the RNA flexibility and RNA-protein/ligand complexes, future breakthroughs may come from integrating deep learning methods with experimental or template information.

Finally, trRosettaRNA2 shows a distinct advantage in predicting the heterogeneous conformations of RNase P RNA, likely stemming from its effective use of diverse SS inputs. Unlike AF3, trRosettaRNA2 produces a conformational ensemble with greater diversity in closer agreement with experimental AFM data. This capability positions trRosettaRNA2 as a promising tool for investigating RNA conformational landscapes, a fundamental and challenging area within RNA biology.

## METHODS

### trRosettaRNA2 algorithm

Different from trRosettaRNA, trRosettaRNA2 employs an end-to-end architecture with three modules: the Secondary Structure (SS) module, the RNAformer module, and the SS-aware structure module.

### SS prior module

The SS prior module generates a base-pairing probability map from the input MSA. The MSA is initially embedded into MSA and pair representations, which are subsequently updated through 8 RNAformer blocks. Finally, a 2D map containing the base-pairing probabilities for all residue pairs is predicted from the updated pair representation.

### RNAformer module

The RNAformer module largely inherits from trRosettaRNA, i.e., interactively updating MSA and pair representations and predicting the 2D geometries using a Res2Net-enhanced Transformer network. The primary distinction lies in the reduced block number: RNAformer in trRosettaRNA2 employs only 12 blocks, down from 48 in trRosettaRNA, as we observed minimal performance impact from this reduction. This module outputs the updated representations, which are subsequently used to generate the full-atom 3D structure by the structure module (described later). Meanwhile, this module also predicts the 2D geometries. Different from trRosettaRNA, trRosettaRNA2 predicts only inter-residue distances and contacts. Inter-residue angles were excluded due to their marginal contribution to trRosettaRNA2’s accuracy and the substantial increase in computational cost for energy minimization.

### SS-aware structure module

The SS-aware structure module generates a 3D structure from the updated representations, assisted by secondary structure (SS) information. This module is primarily driven by Invariant Point Attention (IPA), a mechanism adopted in AlphaFold2 ^9^, which updates the single representation using multi-source attention maps. Specifically, starting with the single representation (the first row of the updated MSA representation) and the pair representation, this module iteratively updates the single representation via IPA and then refines the backbone pose (position and orientation, which are initialized by a Graph Transformer block ^58^) of each nucleotide. This iterative process is repeated 8 times with shared parameters. Finally, side-chain torsion angles are predicted and used to reconstruct full-atom structures. In trRosettaRNA2, the N1(pyrimidine)/N9(purine)-C1’-C4’ frame defines the backbone of each nucleotide, with remaining atom positions reconstructed from predicted torsion angles.

Considering the unique importance of SS for RNA 3D structure, we attempt to integrate SS into the structure module. Compared to the vanilla softmax operation used in self-attention ^59^(Eq. (1)), the SS-aware structure module adopts an SS-weighted softmax. This involves multiplying the base-pair probability map with the attention maps during softmax (i.e., the SS-weighted softmax, see Eq. (2) and Fig. 1C). To prevent over-attenuation of the original attention values, base-pair probabilities are processed using an annealing-smoothing function, with temperature *T* gradually decreases across network blocks (see Fig. S9). Formally, the vanilla attention softmax operation and the SS-weighted softmax operation can be written as:

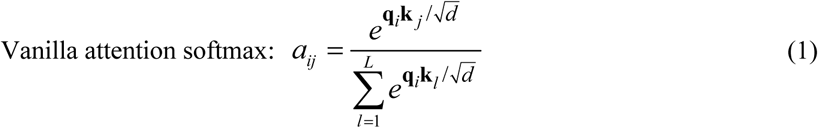

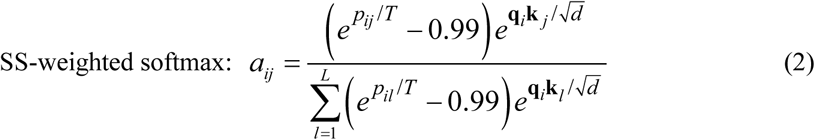

where *aij* is the attention weight assigned by the *i*-th residue to the *j*-th residue; **q***i* and **k***j* refer to the query and key vectors of the *i*-th and *j*-th residues, respectively; *d* represents the dimensionality of the query and key vectors; *L* is the sequence length; *pij* refers to the base-pairing probability between the *i*-th and *j*-th residues; *T* is the temperature factor of the annealing-smoothing function, which shows a base-2 exponential dependence on the network block count.

### Recycling

After each network cycle, the first row of MSA representation, the pair representation, and the distance map extracted from the predicted structure are recycled as input for the next cycle. The whole network undergoes a total of 4 iterations, i.e., with 3 recycling steps.

### Relaxation

Although the training objective of the end-to-end network includes structural violation loss (see *Training procedure*), minor violations can persist in the raw network-generated structures. To address this issue, we implement a rapid relaxation protocol in PyRosetta. This protocol idealizes bond geometries and reduces steric clashes through a fast *MinMover* procedure utilizing the “ref2015_cart” score function within Rosetta. This rapid, yet effective, relaxation step resolves the majority of structural violations, allowing for fast and accurate structure prediction.

### Energy minimization

Beyond the end-to-end prediction, trRosettaRNA2 retains the option for energy minimization using predicted 2D geometries. This procedure is analogous to that employed in trRosettaRNA, except that we exclusively utilize inter-residue distance restraints. Orientation restraints were excluded as their incorporation did not improve trRosettaRNA2’s performance and substantially increased the minimization time. The specific energy function utilized is as follows:

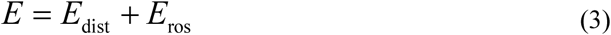

where *E*dist and *E*ros represent the distance-based restraints and Rosetta’s internal energy terms, respectively.

### Training data preparation

#### Training set from PDB

Our primary training dataset was derived from the Protein Data Bank (PDB). We collected all RNA-containing entries released in the PDB before 2024-01 and isolated the RNA components. The multi-chain entries were systematically split and subsequently reassembled into individual chains according to inter-chain base-pairing interactions. The chains shorter than 10 nucleotides were removed. The resulting dataset comprised 10,699 RNA chains, which were used to train both trRosettaRNA2 and trRNA2-SS. To ensure rigorous benchmark evaluations and avoid data leakage, a subset of 8,598 RNAs released before 2022 was designated as the training set for benchmarking.

#### Training set from bpRNA

To facilitate transfer learning for secondary structure prediction in trRNA2-SS, we incorporated data from the bpRNA database. In detail, we collected the bpRNA sequences that are also annotated in the Rfam database, allowing for the direct use of established Rfam MSAs. To reduce redundancy, sequences were filtered at an 80% sequence identity threshold. The final bpRNA training set consists of 14,648 RNA chains.

### Test data preparation

#### Test sets for RNA 3D structure prediction

We use two test sets, TS39 and CASP15, to benchmark trRosettaRNA2’s performance in RNA 3D structure prediction. TS39 was constructed by collecting RNA-only entries in PDB released after 2022-01, splitting and reassembling multi-chain entries according to base-pairing, and removing chains shorter than 20 nucleotides or longer than 400 nucleotides. Redundant sequences within TS39 were removed at an 80% sequence identity cutoff. Finally, RNAs with significant BLASTN hits (E-value < 10) in either the PDB training set (2022- 01 version) or the bpRNA training set were excluded, yielding a final dataset of 39 RNAs. The CASP15 set comprised 10 RNAs from CASP15 experiments, excluding R1189 and R1190 due to their complicated protein interactions. Among these 10 RNAs, 6 were naturally occurring, and 4 were synthetic.

#### Test sets for RNA secondary structure prediction

ArchiveII and PDB27 test sets were used to evaluate trRNA2-SS for RNA secondary structure prediction. ArchiveII, a commonly used benchmark dataset, was filtered to remove RNAs homologous to the training sets (E-value < 10). Additionally, RNAs containing unknown bases (“N”) or longer than 600 nucleotides were removed from ArchiveII, as several compared methods are not equipped to handle such sequences. The final ArchiveII test set comprised 279 RNAs. The second set, PDB27, is a subset derived from TS39, containing 27 RNAs from TS39 that do not have unknown bases (“N”).

For enhanced clarity and overview, a comprehensive summary of all datasets utilized in this work is presented in Table S5.

### Training procedure

#### Pre-training of SS prior module

The SS prior module (i.e., trRNA2-SS) was pre-trained in a transfer-learning manner on the bpRNA and PDB datasets. For the bpRNA samples, the training labels are the base pairs extracted from the dot-bracket notations given by the bpRNA database. For the PDB samples, the labels are the base pairs extracted from the PDB files using DSSR ^60^. The loss function is defined as the cross entropy between the predicted base-pairing probability and the one-hot encoding of true labels. The training was conducted on an A100 40GB GPU, using an Adam optimizer with a learning rate of 0.0002. We trained 3 models with identical configurations. The final trRNA2-SS prediction is the ensemble (average) of the predictions from these 3 models.

#### Training of trRosettaRNA2

The training of trRosettaRNA2 consists of three stages (Table S6). In the first stage, trRosettaRNA2 is primarily trained to produce structures with high accuracy. The loss items include the frame-aligned-position-error (FAPE) loss ℒ_FAPE_, the 2D geometries loss ℒ_geo2D_, the torsion loss ℒ_torsion_, the pLDDT loss ℒ_pLDDT_, and an extra loss item ℒ_Bp_ to optimize the inter-atom distances of the predicted structure according to the native base-pairing information. Specifically, for each native base pair, ℒ_Bp_ forces all the inter-atom distances between these pair of nucleotides in the predicted structure to fall within the range of [1.5 Å, *d* + 3 Å], where *d* is the corresponding inter-atom distance in the experimental structure. In total, the loss function used in the first stage can be written as:

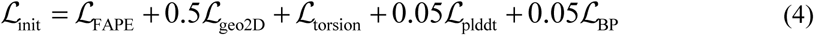

The second stage is the *violation fine-tuning*, which fine-tunes the model to improve the physicochemical plausibility of predicted structures. Specifically, this stage introduces two new loss items. The first item, ℒ_bond_, enforces the length of the O3’-P and P-P bond, as well as the O3’-P- O5’ angle between neighboring residues, to be within ideal ranges, which were statistically derived from the RNAs in the training set. The second is the clash loss ℒ_clash_, which penalizes steric clashes in the predicted structures. In detail, ℒ_clash_ penalizes the inter-atom distances shorter than *min*(*d* − 0.1 Å, 2 Å), where *d* is the corresponding distance in the experimental structure. In total, the loss function used in the second stage can be written as:

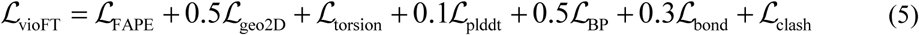

For the first two stages, the training RNAs are clustered based on their sequence identities. During each training epoch, we iterate through all the clusters, sampling one RNA from the current cluster at each step.

The third stage, termed *resampled fine-tuning*, further fine-tunes the model to improve performance on challenging targets. This is achieved by employing a dynamic training sampler that increases the sampling weights for targets exhibiting low lDDT scores. Specifically, each training epoch involves sampling 8000 training examples, where the sampling weight for each RNA is defined as the reciprocal of its lDDT value in the previous epoch. The loss function of this stage is the same as the second stage, i.e.,

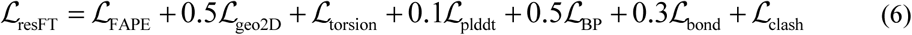

More training details, including learning rates, crop size, and training times, are shown in Table S6.

### Experiment set-up

#### RNA 3D structure prediction

We benchmark trRosettaRNA2 against its predecessor, trRosettaRNA, and other representative deep-learning methods: AlphaFold 3, RhoFold+, DeepFoldRNA, and RoseTTAFoldNA. All methods utilize identical multiple sequence alignments (MSAs) as input. For methods employing secondary structure (SS) input (trRosettaRNA2, trRosettaRNA, and DeepFoldRNA), the specific SS sources are detailed in Table 1 and Fig. 4A. AlphaFold 3 was run using its standalone package with five seeds (0-4), generating five models per seed. The final AlphaFold 3 prediction was selected as the model with the highest pTM score among the 25 generated structures. Due to observed relaxation issues with RhoFold+ for several long RNAs (listed in Table S7), unrelaxed predictions were evaluated in those instances for RhoFold+. All other methods were executed using their publicly available standalone packages with default settings.

### RNA secondary structure prediction

To assess the performance of RNA secondary structure prediction, we benchmark trRNA2-SS against a diverse set of representative methods. This set included traditional energy- and/or evolution-based approaches (EternaFold, RNAfold, RNAstructure, PETfold) as well as advanced deep learning-based methods (RNA-FM, RNA-MSM, UFold, MXfold2, SPOT-RNA, SPOT-RNA2). For methods that utilize MSAs as input (trRNA2-SS, PETfold, SPOT-RNA2, and RNA-MSM), we ensure consistency by providing them with the same MSA inputs. For deep learning-based methods that employ transfer learning, i.e., those pre-trained on secondary structure datasets and finetuned on PDB data (including SPOT-RNA, SPOT-RNA2, and UFold), we evaluate their PDB-finetuned versions.

### Evaluation

To comprehensively assess the performance of trRosettaRNA2, we evaluated its predicted 3D structures using multiple metrics. Among them, root-mean-square deviation (RMSD) and TM-score ^61, 62^ focus on the global topology; the local distance difference test (lDDT) ^63^ to assess local deviations; and interaction network fidelity (INF) ^64^ to measure the accuracy of base-pairing interactions. Unless otherwise specified, the RMSD (based on all atoms) and INF are calculated using the RNA_assessment toolkit ^65^; TM-score is calculated using the RNAalign program ^62^; lDDT is calculated using the following equation:

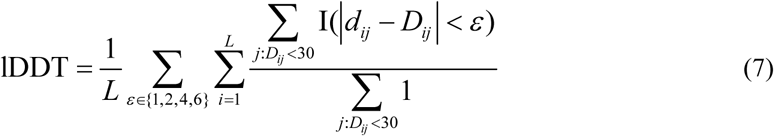

where *L* is the sequence length; *dij* and *Dij* are the C1’ distances between the *i*-th and *j*-th residues in the predicted and experimental structures, respectively; I is the indicator function. To accommodate the typically larger scale of RNA inter-residue distances compared to proteins, we modified the standard lDDT calculation in two aspects. First, the local neighborhood radius used for comparisons increased from 15 Å to 30 Å. Second, the set of distance error thresholds was changed from {0.5 Å, 1 Å, 2 Å, 4 Å} to {1 Å, 2 Å, 4 Å, 6 Å}. Compared to the original formulation, this modified lDDT exhibits a stronger correlation with RMSD (PCC: -0.57 vs. -0.78; Fig. S10).

For the evaluation of secondary structure prediction, we primarily use the Area Under the Precision-Recall Curve (AUPRC), which can provide a direct evaluation of the raw output (i.e., the base-pairing probabilities) generated by various methods.

### Application of trRNA2-SS to RNAcentral sequences

We apply trRNA2-SS to perform high-throughput SS predictions for RNAcentral sequences that lack known SSs. Specifically, we downloaded a list of RNAcentral entries meeting the following criteria: available genomic mapping, no known secondary structure, and the sequence length between 20 and 300 nucleotides. Using this list, we collected the corresponding sequences from our localized RNAcentral database, yielding 588,227 sequences. Redundant sequences were then removed using the CD-HIT program ^66^ with a sequence identity cutoff of 80%. The final set contains 98,008 non-redundant RNA sequences without known SSs. We then used trRNA2-SS to predict the SSs for these sequences. Binary base pairs were derived from the predicted probabilities using a simple cutoff of 0.5.

To facilitate the evaluation of the SS predicted by trRNA2-SS for these RNAcentral sequences, we trained a confidence score (C-Scoress) on 1,000 RNAs randomly sampled from the training set. This score correlates well with the actual F1-score on the test sets (PCC=0.68; Fig. S1). Specifically, C-Scoress can be written as:

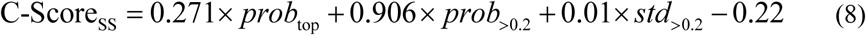

where *prob*top is the average probability of the top *L*/10 predicted base pairs (*L* is sequence length) that have a sequence separation greater than 12; *prob*>0.2 is the average probability of the pairs with probabilities >0.2; *std*>0.2 is the average standard deviation of the probabilities predicted by the 3 individual trRNA2-SS models, calculated only for those base pairs where the final predicted probability is greater than 0.2.

## AVAILABILITY

The web server is available at: https://yanglab.qd.sdu.edu.cn/trRosettaRNA/. The source codes will be available on GitHub soon.

## ACKNOWLEDGEMENTS

This work is supported by the following funding sources: National Natural Science Foundation of China (NSFC T2225007, T2222012, 32430063), Postdoctoral Fellowship Program and China Postdoctoral Science Foundation (BX20240212), and Fundamental Research Funds for the Central Universities.

