## Supplementary Material for "Predicting RNA 3D structure and conformers using a pre-trained secondary structure model and structure-aware attention"

### Supplementary Materials

**Table S1. Comparison between trRosettaRNA2 and DRfold2 on the 34 RNAs.**

| **Method** | **SS predictor** | **RMSD (Å) ↓** | **lDDT ↑** | **TM-score ↑** | **INF ↑** | **clashscore ↓** |
|---|---|---|---|---|---|---|
| trRosettaRNA2 (pyRosetta) | SPOT-RNA | 10.8 | 0.669 | 0.288 | 0.737 | 2.1 |
| | trRNA2-SS | 10.2 | 0.680 | 0.299 | 0.738 | 2.2 |
| | pLDDT top1 | **9.9** | 0.682 | 0.306 | **0.749** | **1.8** |
| DRfold2 (CSOR) | \ | 10.7 | **0.688** | **0.333** | 0.748 | 35.3 |
| DRfold2 (best single model) | \ | 11.0 | 0.653 | 0.313 | \ | \ |

**Table S2. Comparisons on six natural RNAs from CASP15**. For the evaluation of AIchemy_RNA, DF_RNA, and BAKER, we use the same configuration as described in the trRosettaRNA paper. Specifically, we prioritize the evaluation of their submitted models during CASP15, and in cases where a submitted model is unavailable, we replace it with a locally generated model via the corresponding algorithms.

| **Method** | **RMSD (Å)** ↓ | **lDDT** ↑ | **TM-score** ↑ | **INF** ↑ | **clashscore** ↓ |
|---|---|---|---|---|---|
| AIchemy_RNA [H] (RhoFold) | 10.9 | 0.663 | 0.354 | 0.763 | 21.0 |
| RhoFold+ | 11.6 | 0.633 | 0.321 | 0.687 | 32.3 |
| DF_RNA [H] (DeepFoldRNA) | 11.6 | 0.673 | 0.341 | 0.744 | 48.7 |
| BAKER [H] (RoseTTAFoldNA) | 13.8 | 0.613 | 0.302 | 0.734 | 28.9 |
| AlphaFold 3 | <u>9.5</u> | **0.754** | **0.428** | **0.834** | 16.5 |
| Yang-Server[S] (trRosettaRNA) | 12.3 | 0.599 | 0.319 | 0.699 | 26.9 |
| AIchemy_RNA2 [H] | 13.4 | <u>0.718</u> | 0.387 | <u>0.821</u> | <u>12.7</u> |
| trRosettaRNA2 | **9.2** | 0.708 | <u>0.399</u> | 0.791 | **2.6** |

[H]: Human group; [S]: Server group.

**Table S3. Comparisons on four synthetic RNAs from CASP15**.

| Group/Method | RMSD (Å) ↓ | lDDT ↑ | TM-score ↑ | INF ↑ | clashscore ↓ |
|---|---|---|---|---|---|
| AIchemy_RNA[H] (RhoFold) | 40.3 | 0.492 | 0.236 | 0.788 | 10.0 |
| RhoFold+ | 46.3 | 0.309 | 0.168 | 0.212 | 235.8 |
| DF_RNA[H] (DeepFoldRNA) | 54.1 | 0.486 | 0.211 | 0.777 | 63.4 |
| BAKER[H] (RoseTTAFoldNA) | 40.1 | 0.421 | 0.221 | 0.706 | **0.2** |
| AlphaFold 3 | 36.5 | 0.520 | 0.252 | 0.834 | 8.6 |
| Yang-Server[S] (trRosettaRNA) | 39.0 | 0.468 | 0.213 | 0.563 | 165.4 |
| AIchemy_RNA2[H] | **11.3** | **0.753** | **0.692** | **0.884** | 8.7 |
| trRosettaRNA2 | 29.1 | 0.492 | 0.278 | 0.689 | 14.9 |

[H]: Human group; [S]: Server group.

**Table S4. Comparisons of Yang-Server and AF3-Server on 23 targets shorter than 400 nucleotides or with templates**.

| Group | Model | RMSD (Å) ↓ | lDDT ↑ | TM-score ↑ | GDT-TS ↑ | INF ↑ | clashscore ↓ |
|---|---|---|---|---|---|---|---|
| Yang-Server | First model | 10.6 | 0.634 | 0.530 | 0.483 | 0.801 | 3.4 |
| | Best model | **7.8** | 0.663 | **0.573** | **0.518** | 0.826 | **1.5** |
| AF3-server | First model | 10.8 | 0.658 | 0.511 | 0.473 | 0.818 | 14.6 |
| | Best model | 9.8 | **0.676** | 0.535 | 0.494 | **0.829** | 10.6 |

**Table S5. Information of the datasets used in this work.**

| Type | Name | Size | Usage |
|---|---|---|---|
| Training sets | PDB | 10,699 | Training trRNA2-SS and trRosettaRNA2 |
| | bpRNA | 14,648 | Pre-training trRNA2-SS |
| Test sets | TS39 | 39 | Benchmarking trRosettaRNA2 |
| | CASP15 | 10 | |
| | ArchiveII | 279 | Benchmarking trRNA2-SS |
| | PDB27 | 27 | |

**Table S6. Training details of trRosettaRNA2.**

| Parameter | Stage 1: initial training | Stage 2: violation fine-tuning | Stage 3: resampled fine-tuning |
|---|---|---|---|
| Sequence crop size | 256 | 384 | |
| Initial parameters | Random | From stage 1 | From stage 2 |
| Initial learning rate | 0.0002 | 0.00005 | 0.00007 |
| Training sample* | 3,665/4,992 clusters (8,598/10,699 RNAs) | | 8,598/10,699 RNAs |
| Training sampler | By sequence cluster (identical weights for all clusters) | | 8000 samples (weight: reciprocal of the lDDT in the previous epoch) |
| Training times | ~5 days | ~3 days | ~4 days |
| Training resource | 1 A100 40GB GPU | 1 A800 GPU | |

* The numbers preceding the "/" indicate the subset used for benchmarking, which includes RNAs released before 2022, while the numbers following the "/" represent the total number of training RNAs.

**Table S7. Information of the RNAs for which RhoFold+ fails to perform relaxation.**

| Source | ID | Length | Clashscore of RhoFold+ unrelaxed model |
|---|---|---|---|
| CASP15 | R1126 | 363 | 156.5 |
| | R1136 | 374 | 186.5 |
| | R1138 | 720 | 137.5 |
| CASP16 | R1248 | 407 | 579.0 |
| | R1283 | 580 | 432.2 |
| | R1286 | 526 | 222.2 |

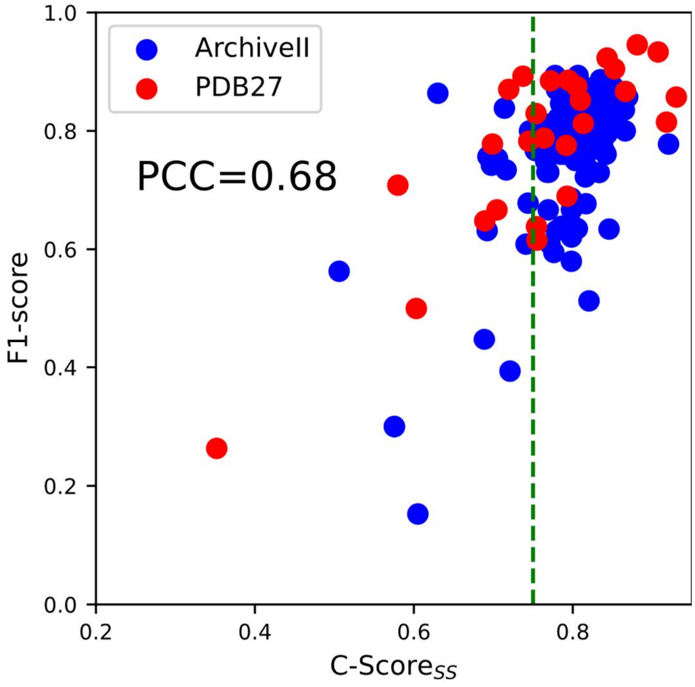


**Fig. S1. Relationship between F1-Score and C-Score$_{SS}$ on two test sets.** For 91.1% of the targets exhibiting a C-Score$_{SS}$ greater than 0.75 (green dashed line), the corresponding F1-scores also exceeded 0.75.

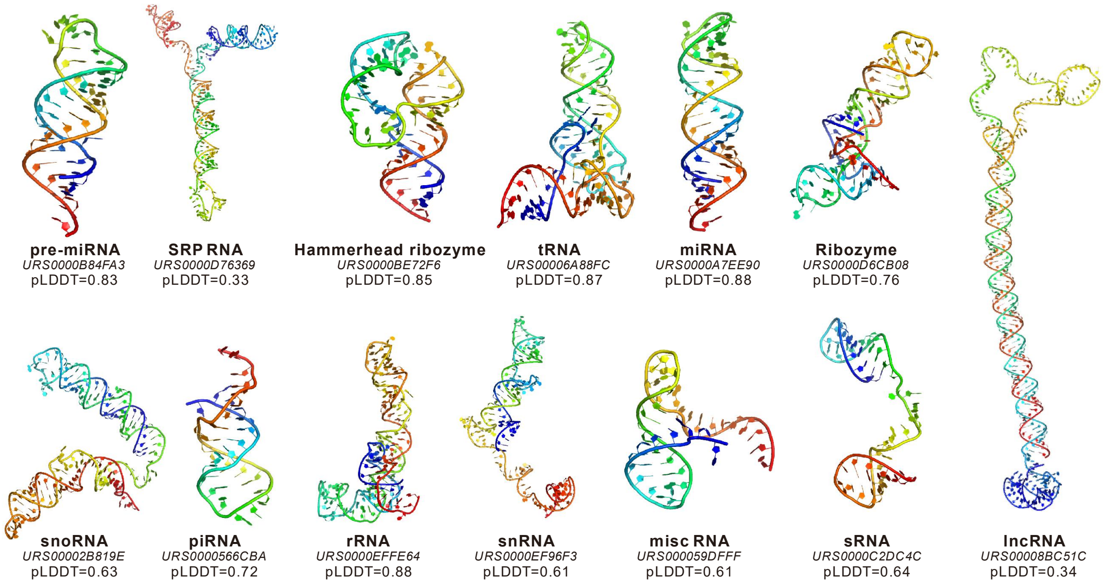


**Fig. S2. 3D structures predicted by trRosettaRNA2 for the representative RNAs from RNAcentral.**

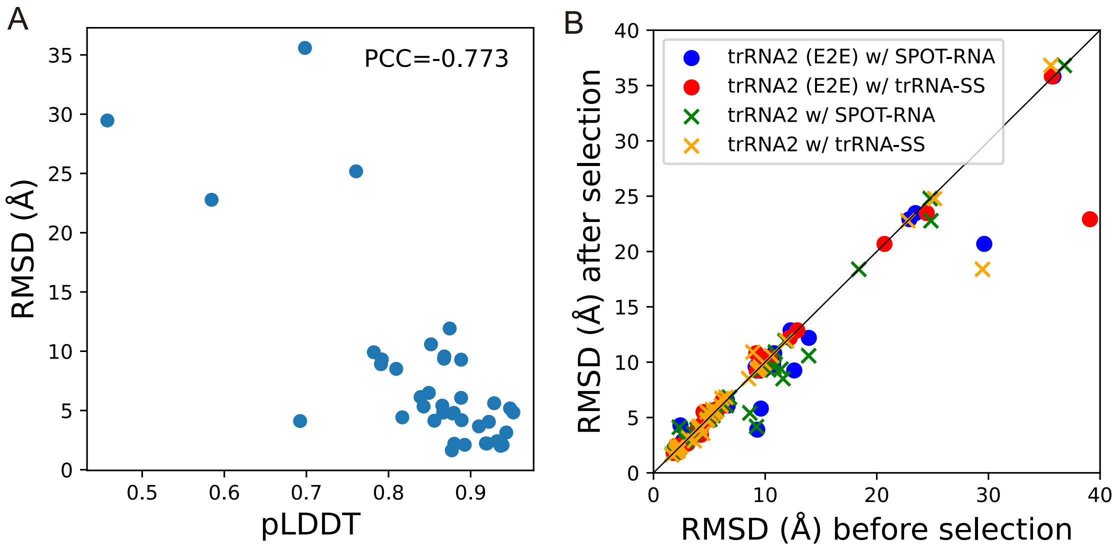


**Fig. S3. Relationship between pLDDT and RMSD on the TS39 test set (A) and its application to model selection (B).**

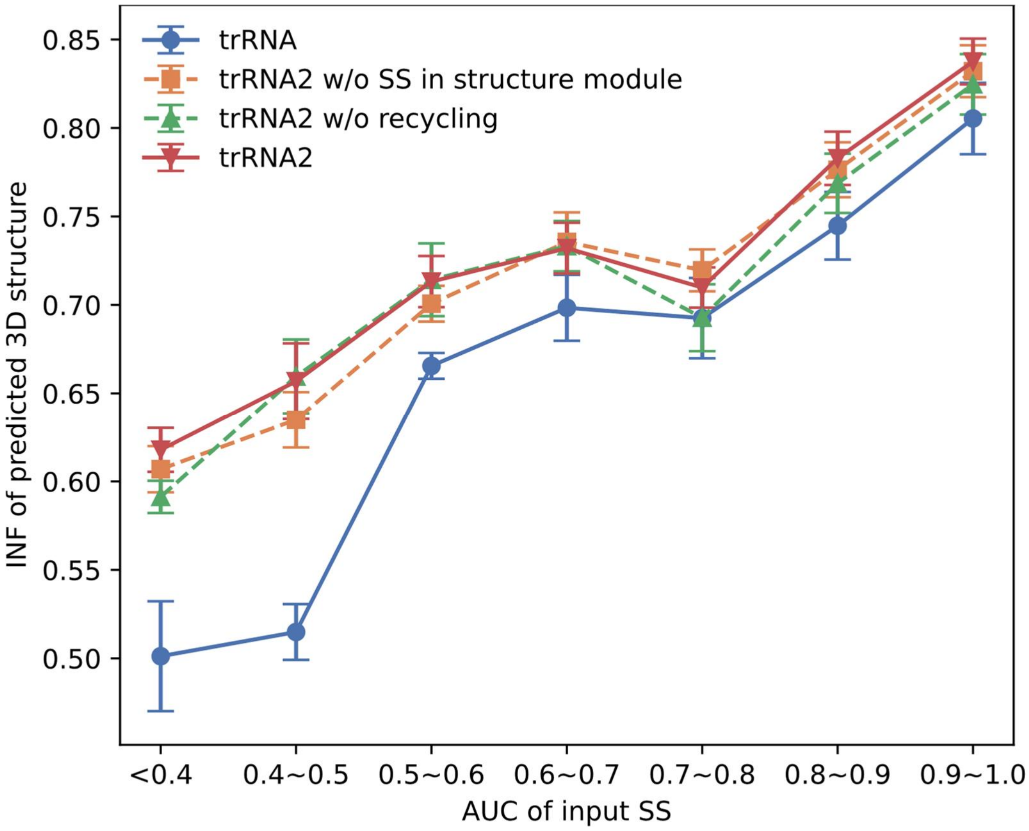


**Fig. S4. Relationship between INF of predicted 3D structure and AUC of input SS.** The error bars indicate one-fifth of the standard deviation. “trRNA2” refers to trRosettaRNA2 (E2E) here.

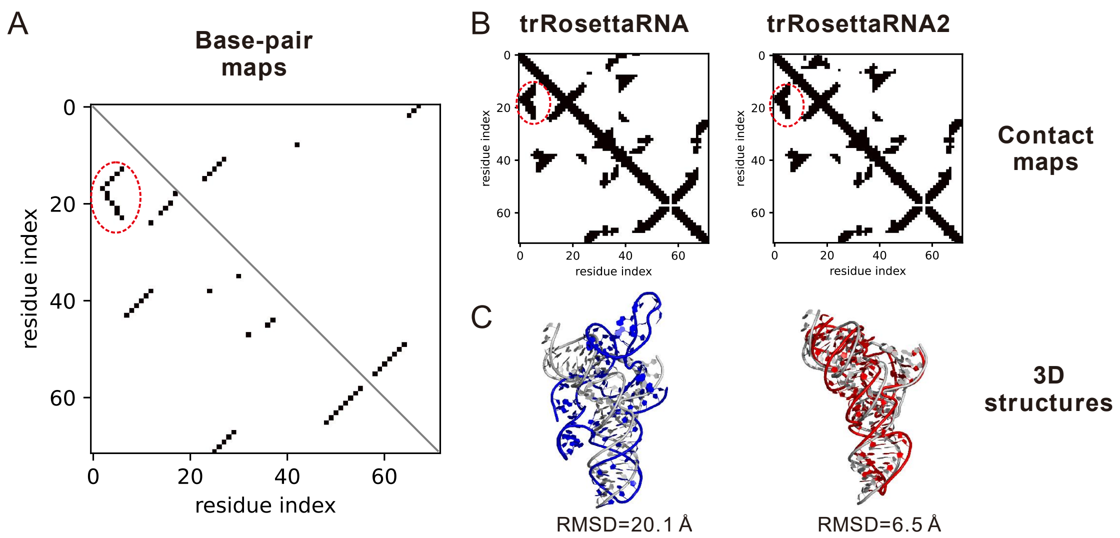


**Fig. S5. trRosettaRNA vs. trRosettaRNA2 on an example RNA (PDB ID: 8YDC) with SPOT-RNA input.** (A) Comparison of the base-pair map predicted by SPOT-RNA (upper right) and that extracted from experimental 3D structure (lower left). (B) Comparison of the inter-residue contact maps predicted by trRosettaRNA/trRosettaRNA2 (upper right) against the contact map derived from the experimental 3D structure (lower left). (C) Superposition of the 3D structures predicted by trRosettaRNA (blue) and trRosettaRNA2 (red) onto the experimental structure (gray). The red dashed circle in (A) and (B) highlight the interactions corresponding to the triple helix.

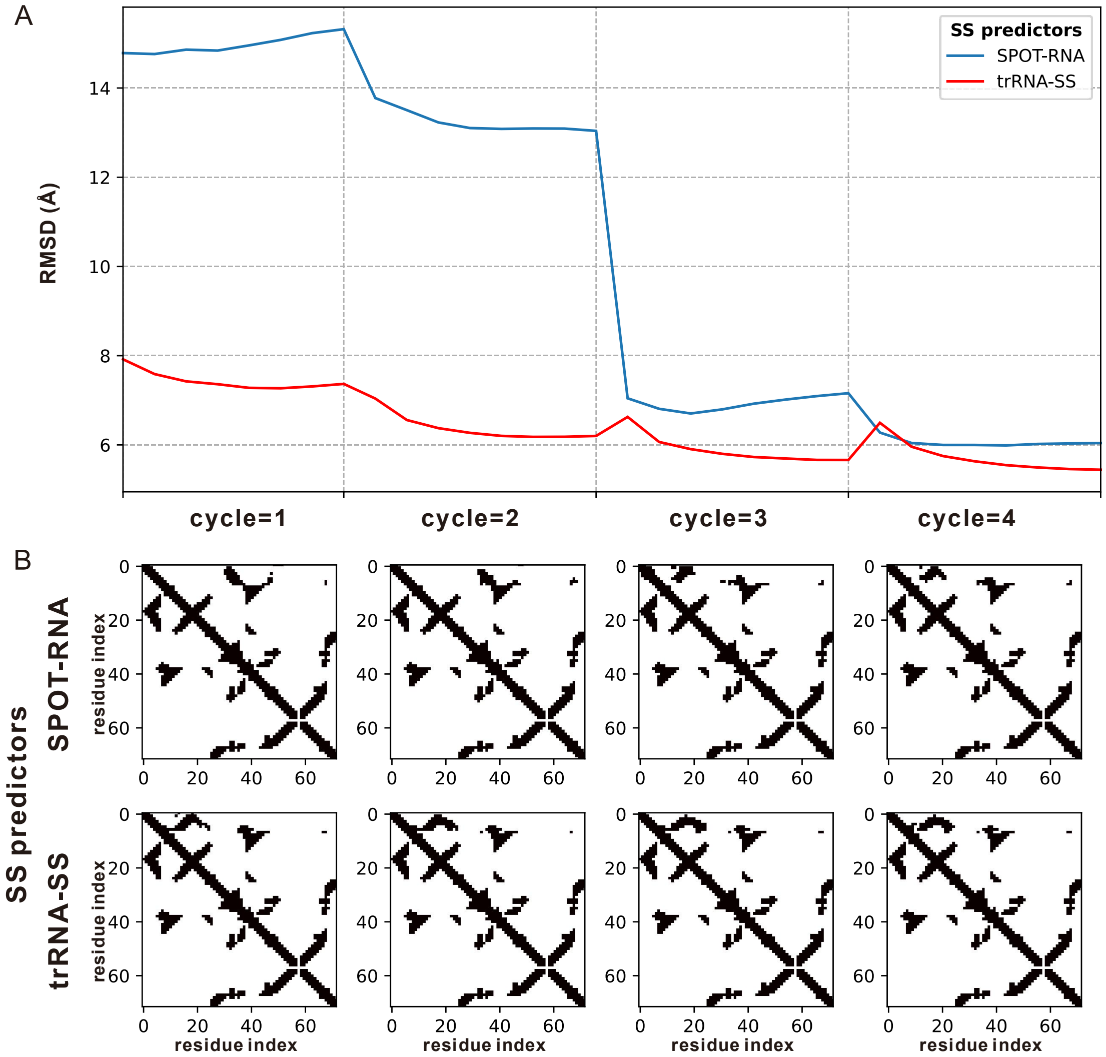


**Fig. S6. The impact of recycling on the prediction of the example RNA (PDB ID: 8YDC).** (A) Relationship between RMSD and number of cycles for trRNA2 using different SS predictors. (B) Comparison of inter-residue contact maps generated at different cycles and using different SS predictors for input. Within each 2D plot in panel (B), the predicted contact map is displayed in the upper right corner, juxtaposed against the experimentally derived contact map shown in the lower left corner.

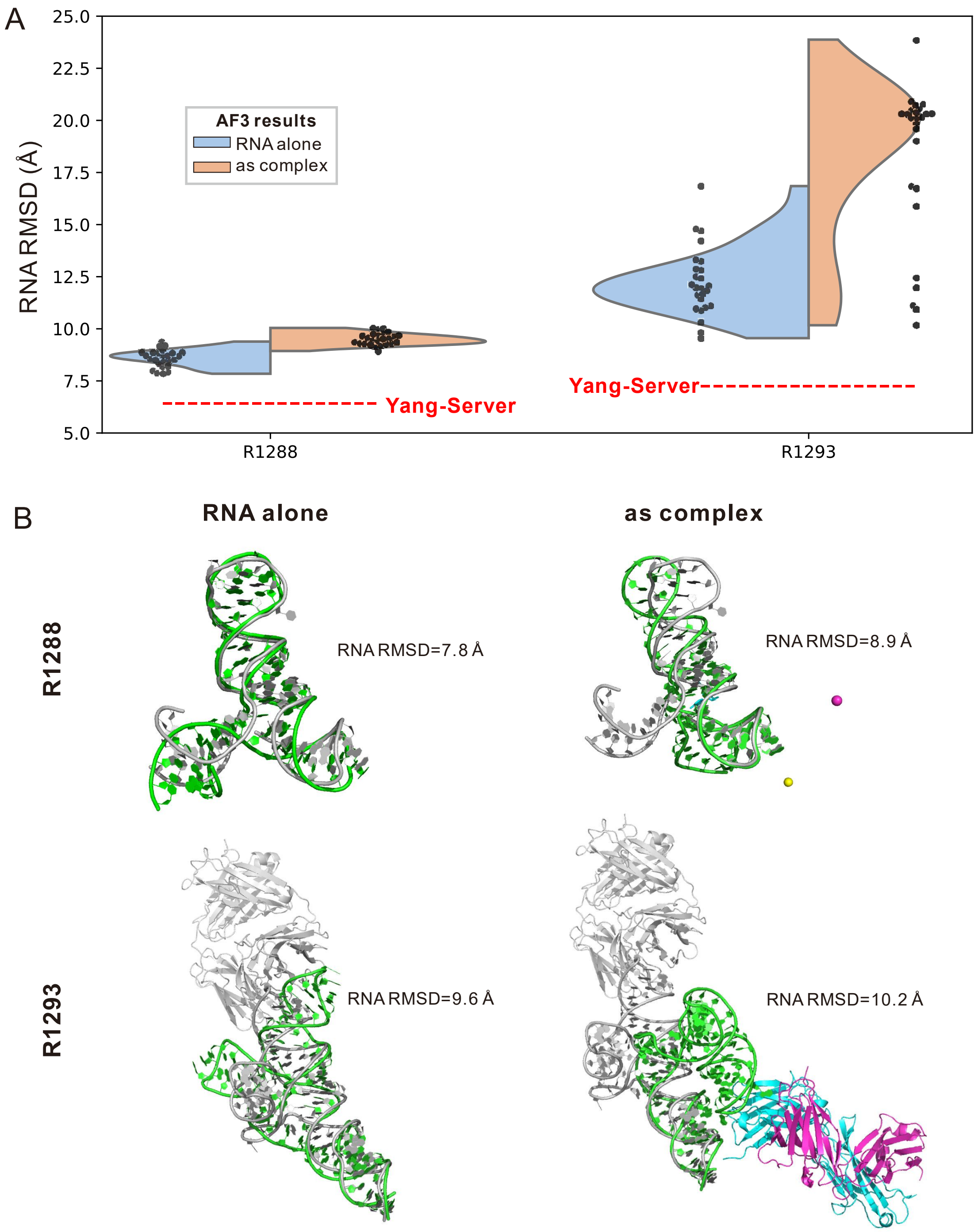


**Fig. S7. Evaluating AF3 predictions for CASP16 hybrid targets R1288 and R1293.** AF3 predictions were generated using the local AlphaFold 3 package with a fixed seed set (0-4), producing 5 models per seed, totaling 25 models for each target. (A) RNA RMSD distribution of the AF3 models when modeling the RNA component alone versus modeling the entire complex. (B) Superimposition of the best AF3 models (RNA in green; proteins or ligands in other colors) onto the experimental structures (in gray).

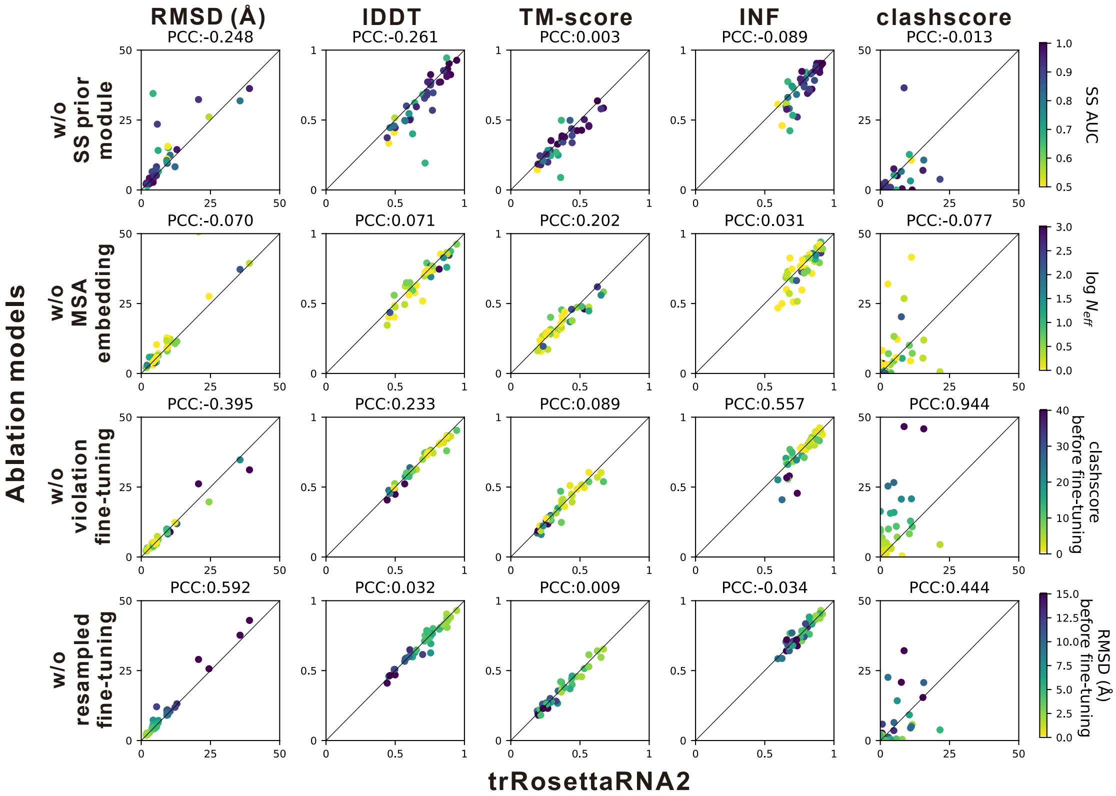


**Fig. S8. Head-to-head comparisons between trRosettaRNA2 and ablation models.** The PCC value above each subplot indicates the correlation between the performance degradation and the quality metric associated with the ablated component. E.g., the second subplot in the first row shows a PCC of -0.261, indicating that the lDDT degradation after removing the SS prior module has a slight negative correlation with the quality (AUC) of SS generated by the SS prior module. This suggests that even for targets where the SS prior module generates low-quality prior information, the removal still tends to worsen the performance.

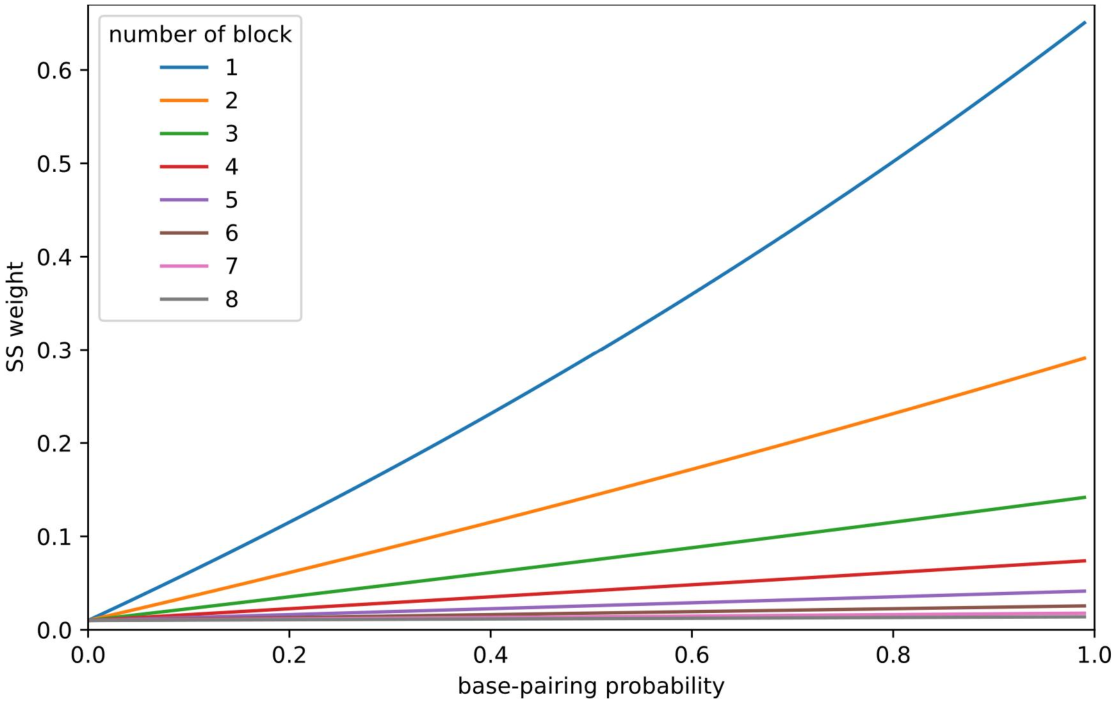


**Fig. S9. Relationship between the SS weight used in the structure module and the original base-pairing probability.** For example, for the first block, the SS weight derived from the highest probability is around 0.6, which is 60 times greater than the weight from the lowest probability. For the last block, the weights are smoothed, with values around 0.01 across the entire range of probabilities.

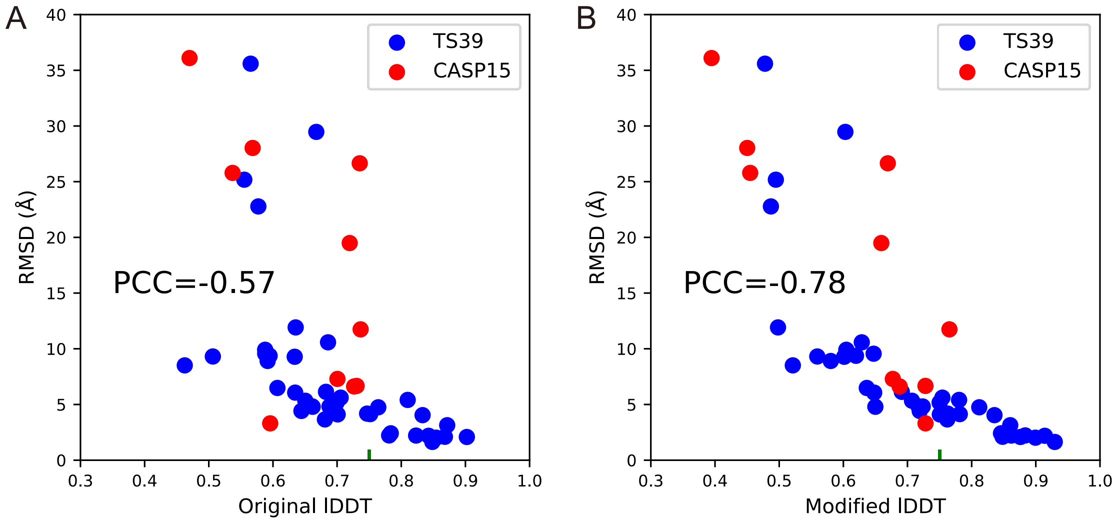


**Fig. S10. Relationship between RMSD and lDDT before and after modification.**
